# Cell-type specific epigenetic clocks to quantify biological age at cell-type resolution

**DOI:** 10.1101/2024.07.30.605833

**Authors:** Huige Tong, Xiaolong Guo, Macsue Jacques, Qi Luo, Nir Eynon, Andrew E. Teschendorff

**Author notes:** Equal contribution (Huige Tong, Xiaolong Guo). Corresponding author: Andrew E. Teschendorff.

## Abstract

The ability to accurately quantify biological age could help monitor and control healthy aging. Epigenetic clocks have emerged as promising tools for estimating biological age, yet they have been developed from heterogeneous bulk tissues, and are thus composites of two aging processes, one reflecting the change of cell-type composition with age and another reflecting the aging of individual cell-types. There is thus a need to dissect and quantify these two components of epigenetic clocks, and to develop epigenetic clocks that can yield biological age estimates at cell-type resolution. Here we demonstrate that in blood and brain, approximately 39% and 12% of an epigenetic clock’s accuracy is driven by underlying shifts in lymphocyte and neuronal subsets, respectively. Using brain and liver tissue as prototypes, we build and validate neuron and hepatocyte specific DNA methylation clocks, and demonstrate that these cell-type specific clocks yield improved estimates of chronological age in the corresponding cell and tissue-types. We find that neuron and glia specific clocks display biological age acceleration in Alzheimer’s Disease with the effect being strongest for glia in the temporal lobe. Moreover, CpGs from these clocks display a small but significant overlap with the causal DamAge-clock, mapping to key genes implicated in neurodegeneration. The hepatocyte clock is found accelerated in liver under various pathological conditions. In contrast, non-cell-type specific clocks do not display biological age-acceleration, or only do so marginally. In summary, this work highlights the importance of dissecting epigenetic clocks and quantifying biological age at cell-type resolution.

## Introduction

Individuals of the same chronological age may display marked differences in health and future disease risk [1]. These differences are attributed to variations in biological age, and whilst a number of biomarkers for biological age have been developed, these are mostly measured in, or restricted to, easily accessible tissues like blood [1–4]. More recently, epigenetic clocks based on DNA methylation (DNAm) marks [5–7] have emerged as powerful tools to predict chronological age, and to some extent also biological age [3, 8–16]. However, many challenges remain that limit the practical utility and interpretation of these epigenetic clocks [9, 10, 17].

Prominent among these challenges is cell-type heterogeneity (CTH) [18, 19]: most if not all epigenetic clocks have been developed or trained from heterogeneous bulk-tissues that are composed of many different cell-types. This CTH has two undesirable effects, as recently stressed by others [20–22]. First, it can distort biological age estimates by underlying variations in cell-type fractions. As a concrete example, whole blood, as well as solid tissues with a significant immune-cell infiltration, are all characterized by an age-related shift from naïve to mature T-cell fractions [23, 24] [25, 26], which as shown by Jonkman et al can confound biological age estimates in immune cell subtypes [23]. Whole blood tissue is also subject to an age-related skew towards the myeloid lineage [27]. Another example is brain, with early studies reporting correlations of an accelerated Horvath-clock[28] with neurodegenerative features (e.g. neurotic plaques or amyloid load) [29, 30] in the prefrontal cortex, or with genetic variants associated with neurodegenerative diseases [31–33], but with more recent studies attributing such age-accelerations to changes in underlying cell-type proportions [34, 35] [36]. The second undesirable effect of CTH is that, even if we adjust for it *a-posteriori*, as it is often done in blood tissue by adjusting epigenetic clock values for variations in estimated immune cell fractions [28], the ensuing epigenetic clock estimates are still averages over all the distinct cell-types in the tissue. Thus, standard *a-posteriori* adjustment for cell-type fractions does not inform us about the biological aging of individual cell-types. This is a serious limitation for understanding which cell-types age faster and contribute most to age-related diseases, or for identifying the cell-types that respond best to cellular rejuvenation and other healthy-age promoting interventions [37–42].

Thus, current epigenetic clocks are composites, reflecting a mixture of two underlying age-related processes: an extrinsic one reflecting age-related variations of a tissue’s cell-type composition, and an intrinsic one, reflecting age-related DNAm changes in the individual cell-types of the tissue, the latter being itself a composite process, as each cell-type may age at a different rate. Importantly, the relative contribution of extrinsic and intrinsic aging processes to an epigenetic clock’s accuracy has not yet been quantified, and the only attempts so far at quantifying biological age at cell-type resolution have come from studies that have applied epigenetic clocks derived from bulk-tissue (e.g. liver) to sparse and unreliable single-cell DNAm data (e.g. hepatocytes) [43].

Here we advance the epigenetic clock field in two ways. First, we dissect epigenetic clocks into their extrinsic and intrinsic components, rigorously quantifying these aging components in two of the most frequently profiled tissues (whole blood and brain), demonstrating that extrinsic aging can account for a substantial proportion of an epigenetic clock’s predictive accuracy. Second, we present a computational strategy for building cell-type specific epigenetic clocks that can yield biological age estimates at cell-type resolution. This strategy explicitly adjusts for CTH whilst training the clock, leveraging recent state-of-the-art cell-type deconvolution methods [24, 44] to identify cell-type specific age-associated DNAm changes [45]. As such this is one of the first studies to characterize age-associated DNAm changes at cell-type resolution. We illustrate our strategy in brain and liver tissue, demonstrating how cell-type specific epigenetic clocks from these tissues can improve tissue-specific estimation of chronological and biological age.

## Results

### Dissecting and quantifying the two components of epigenetic clocks

Epigenetic clocks are composites of two underlying age-related processes, an extrinsic one that reflects age-associated variations in cell-type composition and an intrinsic one that captures age-associated DNAm changes in individual cell-types (**Fig.1a**). Thus, the predictive accuracy (R^2^ value) of a clock can be composed as the sum of an extrinsic and intrinsic accuracy, so that the extrinsic R^2^ value can be estimated from the predictive accuracies of two clocks: one adjusted for CTH, and another which is not (**Fig.1a**). To quantify the relative rates of extrinsic and intrinsic aging, we assembled a large collection of whole blood Illumina DNAm datasets, using two of the largest cohorts to train Elastic Net age-predictors unadjusted and adjusted for CTH, respectively (**Methods**). Adjustment for CTH was performed at the resolution of 12 immune cell subtypes using a validated DNAm reference matrix encompassing naïve and memory B-cells, naïve and memory CD4T-cells, naïve and memory CD8T-cells, T-regulatory cells, natural-killer cells, monocytes, neutrophils, eosinophils and basophils [24] (**Methods**). We also built an elastic net clock adjusting for 9 immune-cell subtypes, by merging together the estimated fractions of the naïve and memory lymphocyte subsets, allowing us to assess the importance of the known age-related shift from naïve to mature T-cell subsets in driving an epigenetic clock’s accuracy (**Methods**) [24]. We then estimated R^2^ values of the 3 clocks in 11 independent whole blood DNAm datasets (**Methods**). This revealed that the unadjusted clock achieved substantially higher R^2^ values than the two adjusted clocks, specially when compared to the clock adjusted for all 12 immune cell types (**Fig.1b, SI figs.S1-2**). We estimated that the intrinsic aging component accounts for approximately 61% of the predictive accuracy of the unadjusted clock (**Fig.1b**), which means that extrinsic aging accounts for a substantial fraction of a clock’s accuracy (39%). Of note, not adjusting for relative shifts in naïve to memory subsets, we would have over-and-under estimated the intrinsic and extrinsic aging components to be approximately 84% and 16%, respectively (**Fig.1b**). By assembling a large collection of brain DNAm datasets (**Methods, SI table S1**) and adjusting for 7 brain cell-type fractions (GABAergic excitatory neurons, glutamatergic inhibitory neurons, astrocytes, microglia, oligodendrocytes, endothelial cells and stromal cells), as estimated with the HiBED algorithm [46], we verified that the intrinsic component of aging in brain is around 88%, higher than the estimate obtained in blood (**Fig.1c, SI fig.S3**). To see which brain cell-types are driving the extrinsic aging component, we performed a fixed and random effects meta-analysis over 13 Illumina DNAm brain datasets, encompassing over 2,000 samples (**SI table S1**), which revealed an age-associated increase of excitatory neurons, whilst inhibitory neurons and astrocytes decreased (**Fig.2**). Thus, this age-related shift in neuronal subtypes accounts for the 12% extrinsic aging component. Of note, these findings are broadly consistent with recent single-nucleus DNAm studies reporting reduced inhibitory neuron fractions in the brain of aged donors [47, 48]. Overall, these results obtained in blood and brain tissue highlight the critical importance of adjusting for CTH when building epigenetic clocks.

**Figure-1:**
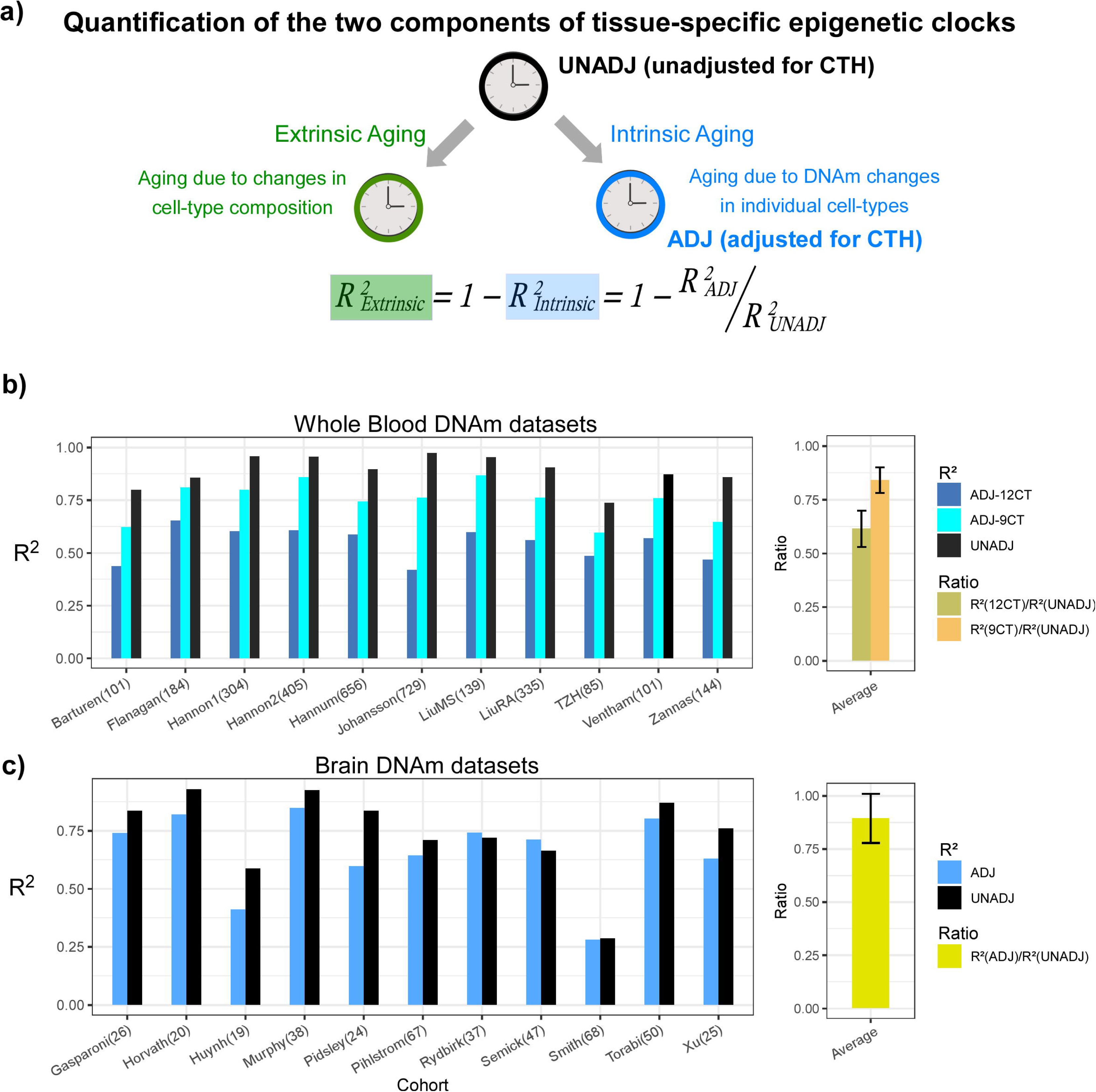
Quantification of intrinsic and extrinsic aging. **a)** Graphical depiction of how the predictive accuracy of an epigenetic clock derived from bulk-tissue can be decomposed in terms of an extrinsic and intrinsic aging process, as shown. The accuracy, measured by the R^2^ value, of a clock reflecting the extrinsic process can be estimated from the given formula, if one can estimate the accuracies of the full unadjusted clock and the one fully adjusted for cell-type heterogeneity (CTH), the latter reflecting the intrinsic aging of the individual cell-types. **b)** Barplots compare the R^2^ values of the unadjusted clock, and clocks adjusted for 12 and 9 immune cell types, in each of 11 whole blood cohorts. Right panel displays the average and standard deviation of the two indicated ratios across the 11 cohorts. These ratios measure the fraction of the unadjusted clock’s accuracy that can be attributed to intrinsic aging. **c)** As b) but for brain DNAm datasets, where the adjusted clock was adjusted for 7 brain cell-types.

**Figure-2:**
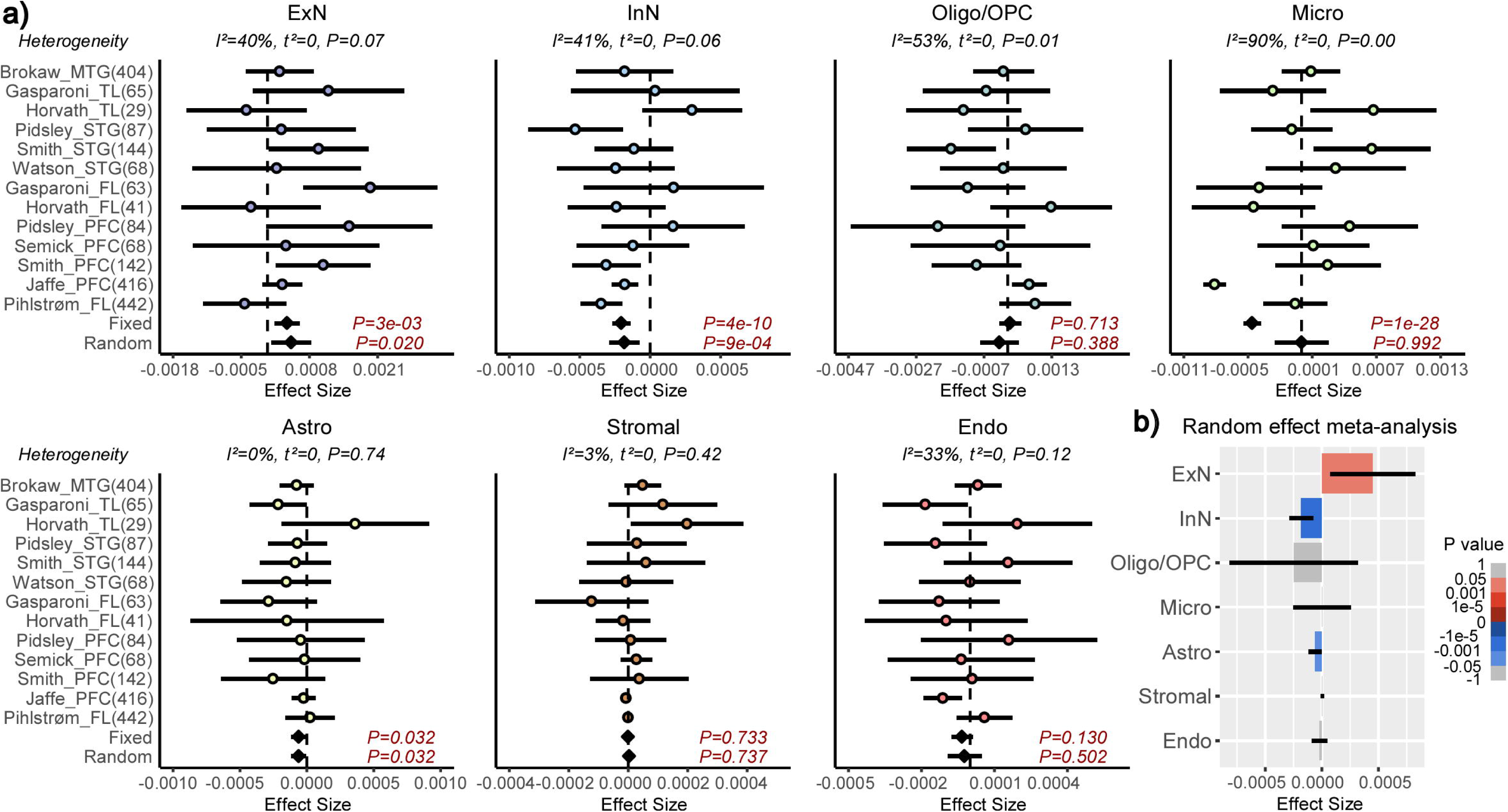
Meta-analysis of brain cell-type fractions with age. **a)** Forest plots of associations between brain cell-type fractions and age, for 7 brain cell-types (ExN=excitatory neurons, InN=inhibitory neurons, Oligo/OPC=oligodendrocytes/oligo precursor cells, Micro=microglia, Astro=astrocytes, Stromal, Endo=endothelial) across 13 independent DNAm datasets. Number of samples in each cohort is given in brackets. P-values from a Fixed and Random Effects models are also given. Heterogeneity statistics and P-values of heterogeneity are given above each plot. **b)** Effect sizes and P-values from the Random Effects meta-analysis model.

### A computational strategy to build cell-type specific epigenetic clocks

In order to dissect the intrinsic aging component, we next devised a computational strategy to build cell-type specific epigenetic clocks (**Fig.3a**). After estimation of the cell-type fractions in a given bulk-tissue DNAm dataset, we apply the CellDMC algorithm [45] to a training subset of the data to identify age-associated differentially methylated cytosines in each cell-type of interest (age-DMCTs) (**Fig.3a**). CellDMC uses the estimated cell-type fractions as weights in a model that expresses a CpG’s DNAm profile across samples as a corresponding weighted mixture of latent phenotype-dependent DNAm profiles. In the present context, the phenotype is age and the described model includes interaction terms between age and the cell-type fractions (**Methods**). These interaction terms allow cell-type specific age-DMCTs to be found (see **Fig.3a** for a hypothetical example). Assuming there is a substantial number of age-DMCTs for a given cell-type, we then apply an Elastic Net Regression model [49] to train predictive models for chronological age parameterized by a penalty parameter (**Fig.3b**). This training is restricted to age-DMCTs of the given cell-type and can be performed either on the residuals of the DNAm data matrix, obtained by regressing out the effect of cell-type composition, or on the unadjusted DNAm data matrix (**Methods**). In the former case, we define the clocks as being “intrinsic”, otherwise they are “semi-intrinsic”. The best performing models in a blinded model-selection set define our cell-type specific intrinsic and semi-intrinsic epigenetic clocks, one for each cell-type (**Fig.3b**). These clocks are then validated in independent DNAm datasets.

**Figure-3:**
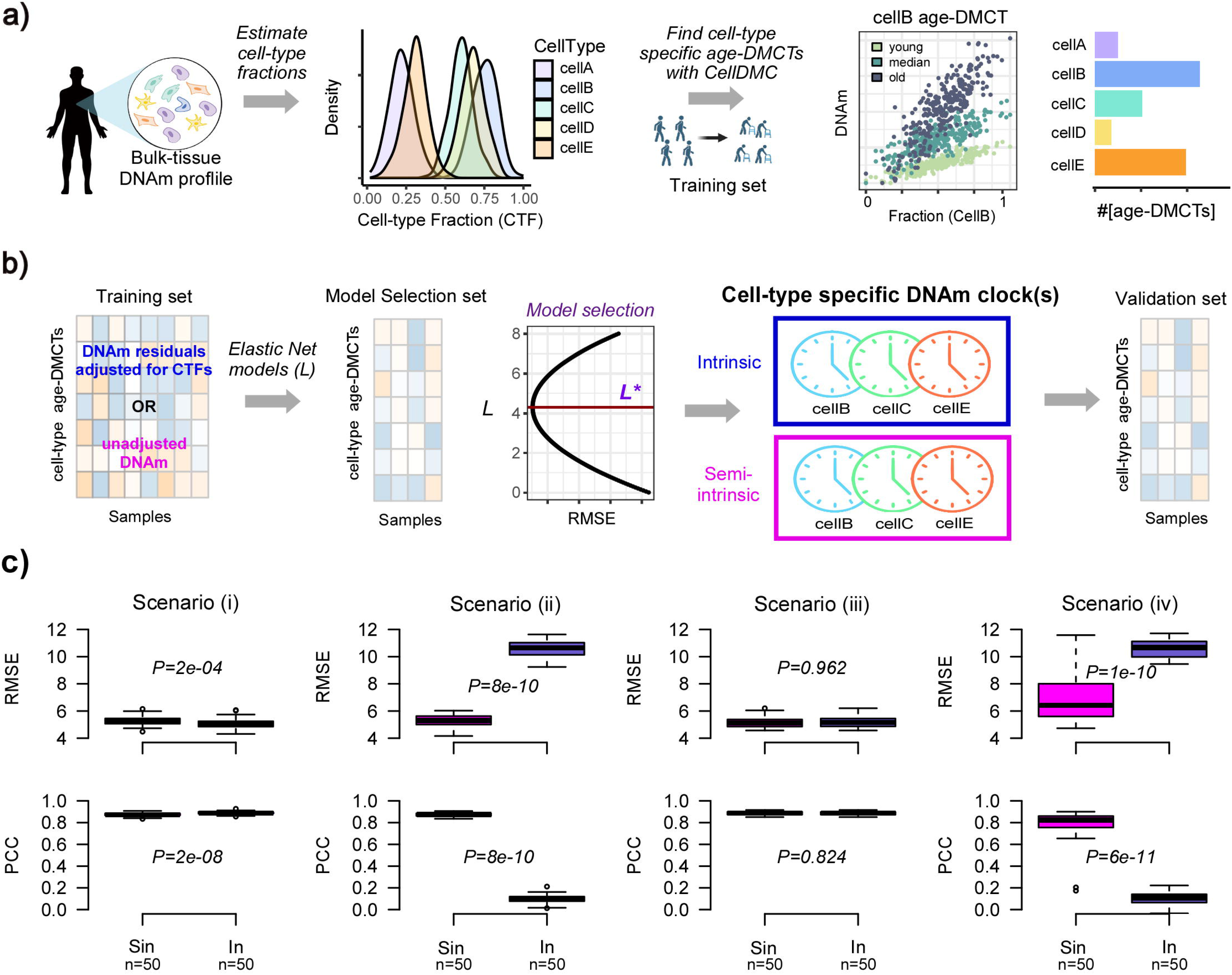
Computational strategy for building cell-type specific epigenetic clocks. **a)** Given a tissue-type of interest with DNAm profiles measured over a relatively large number of samples, we first estimate the underlying cell-type proportions in each sample using an existing tissue-specific DNAm reference matrix. Density plots depict the distribution of cell-type fractions across all samples. Next, using a sufficiently large training set of samples encompassing a relatively wide age-range, we apply the CellDMC algorithm to infer age-associated DNAm changes in each of the underlying cell-types (age-DMCTs). Barplot depicts the number of age-DMCTs in each cell-type. **b)** The construction of cell-type specific clocks then proceeds by restricting to age-DMCTs of one cell-type: an intrinsic clock is built by adjusting the DNAm training dataset for variations in cell-type fractions (CTFs), defining a matrix of DNAm residuals. Alternatively, the training over the age-DMCTs can be done on the DNAm data matrix without adjustment for CTFs which will result in a “semi-intrinsic” clock. In either case, Elastic Net models are learned for each choice of a penalty parameter *L*, and the optimal model *L\** is selected based on the best generalization performance (smallest root mean square error (RMSE)) obtained in a blinded model selection set. This optimal model then defines the corresponding cell-type specific DNAm-clock. This procedure can be done for each cell-type separately, assuming that sufficient numbers of age-DMCTs in that cell-type can be identified. Once the cell-type specific clocks are built, these are then validated in independent DNAm datasets. **c)** Top row: boxplots display the root mean square error (RMSE) between predicted and true ages, for semi-intrinsic (Sin) and intrinsic (In) clocks, as assessed in 50 simulated validation sets of 200 mixtures for four different scenarios. Cell-type specific clocks were constructed from 50 simulated training sets of 200 mixtures (mixing together 3 cell-types that we call granulocytes, monocytes and lymphocytes) with age-DMCTs occurring only in one cell-type (lymphocytes). In scenarios (i) and (iii), no cell-type fractions change with age, in scenarios (ii) and (iv) the lymphocyte fraction changes with age. In scenarios (i) and (ii), age-DMCTs do not discriminate immune-cell types from each other, in scenarios (iii) and (iv) age-DMCTs discriminate lymphocytes from granulocytes and monocytes. P-values from a two-tailed paired Wilcoxon test are given. Bottom row: as top-row but now displaying the PCC of predicted vs true age.

The reasoning behind defining separate intrinsic and semi-intrinsic clocks can be understood with simulation models. We generated simulated DNAm mixtures of 3 underlying cell-types (granulocytes, monocytes and lymphocytes), and in which age-DMCTs are introduced into only one cell-type (lymphocytes) under four different scenarios (**Methods**): (i) age-DMCTs do not map to cell-type specific CpGs and no cell-type fractions change with age, (ii) as (i) but with the lymphocyte fraction changing with age, (iii) age-DMCTs discriminate lymphocytes from the other two cell-types and no cell-type fraction changes with age, (iv) as (iii) but with the lymphocyte fraction changing with age. For each scenario, we generated training and validation sets, deriving lymphocyte specific intrinsic and semi-intrinsic clocks, subsequently validating them in the independent test sets. Interestingly, whilst the intrinsic clock displayed marginally better performance in the scenarios where cell-type fractions did not change with age, the semi-intrinsic clock was substantially better in cases where the lymphocyte fraction did change with age (**Fig.3c**). Thus, the use of residuals in scenarios where a given cell-type fraction changes with age can dilute out the biological signal in that cell-type, and under these circumstances, the cell-type specific semi-intrinsic clock would be preferable.

### Construction and validation of neuron-specific epigenetic clocks

Next, we applied this strategy to the case of brain tissue. For training the clocks, we used one of the largest collections of prefrontal cortex (PFC) samples profiled with Illumina 450k technology [50], encompassing 416 samples with a wide age range (18-97 years) (**SI table S1, Fig.4a**). Although the HiBED algorithm [46] inferred fractions for 7 brain cell-types (**SI fig.S4a**), for subsequent CellDMC analysis, power considerations [45, 51] required us to collapse the inferred fractions to 3 broad cell-types: neurons, glia and endothelial/stromal (**Fig.4a**). Applying CellDMC with the estimated fractions, disease status and sex as covariates, resulted in 16,814 neuron-specific and 11,157 glia-specific age-DMCTs (**Fig.4a, Methods**). Visualization of representative age-DMCTs confirmed the cell-type specific nature of their age-associated DNAm changes (**Fig.4b**). Interestingly, neuron and glia age-DMCTs displayed a weak marginal co-localization with the cell-type specific CpGs in the brain DNAm reference matrix (**SI fig.S5**). Focusing on the neuron-DMCTs and adjusting the DNAm data for variations in cell-type fractions, we next applied the Elastic Net regression framework to learn different models specified by a penalty parameter. For model (i.e. optimal parameter) selection, we used a completely independent Illumina DNAm dataset from Philstrom et al [52] which included 67 PFC control samples with an age range 49-100 years (**Fig.4c, SI table S1**). We declared the model achieving best generalization performance in the Philstrom dataset, as defining our neuron-specific intrinsic (“Neu-In”) clock (**Fig.4c, SI fig.S4b, SI table S2**). This neuron-specific clock was further validated in 5 independent datasets of sorted neuron samples, encompassing a total of 143 samples (**Fig.4d, SI fig.S4c, SI table S1**). An analogous neuron “semi-intrinsic” clock (“Neu-Sin”) was obtained by applying the same training procedure to the same age-DMCTs in the Jaffe et al DNAm data but now without adjusting the DNAm data for variations in cell-type fractions (**Methods, Fig.4d, SI fig.S4d, SI table S3**). To assess specificity of the clocks to neurons and brain tissue, we assembled a large collection of Illumina DNAm datasets encompassing other brain cell-types (glia), other sorted non-brain cell-types and bulk tissue-types (**SI table S1, Methods & Supplementary Methods**). As a further control, we also benchmarked our Neu-In/Sin clocks to two other clocks: a clock obtained by running the Elastic Net on all CpGs but adjusting the DNAm data for cell-type fractions (**Methods, Fig.4d, SI fig.S4e, SI table S4**), thus defining a brain, but not neuron-specific, clock (“BrainClock”), and the multi-tissue Horvath clock. We restricted the assessment of all clocks to control samples only, to avoid potential confounding by disease. First, we observed that the neuron intrinsic and semi-intrinsic clocks also displayed good correlations with chronological age in sorted glia datasets, although the largest of these glia datasets displayed a significantly weaker association, suggesting that our neuron-clocks are indeed neuron-specific (**Fig.4d**). In line with this, a multivariate linear regression of correlation strength against cell-type (neuron vs glia), weighted by cohort size, revealed a significantly higher correlation in neurons compared to glia for the neuron-intrinsic clock but not for the other 3 clocks (**SI fig.S6a**). When assessed between brain and non-brain tissue/cell-types, we observed that the two neuron-specific clocks as well as the brain-specific clock achieved better predictions of chronological age in the brain-tissue datasets compared to non-brain ones, whilst, as expected for a multi-tissue clock, Horvath’s clock performed similarly between brain and non-brain tissues (**Fig.4e-f**). Importantly, and despite the relatively small number of sorted neuron datasets, the neuron-specific clocks also displayed stronger correlations with age in the sorted neuron datasets compared to the brain-specific clock (**SI fig.S6b**). Thus, overall, our data support the view that the neuron-specific clocks are indeed optimized to predict chronological age in brain cell subtypes and that they are less confounded by changes in cell-type composition compared to e.g. the brain-specific clock.

**Figure-4:**
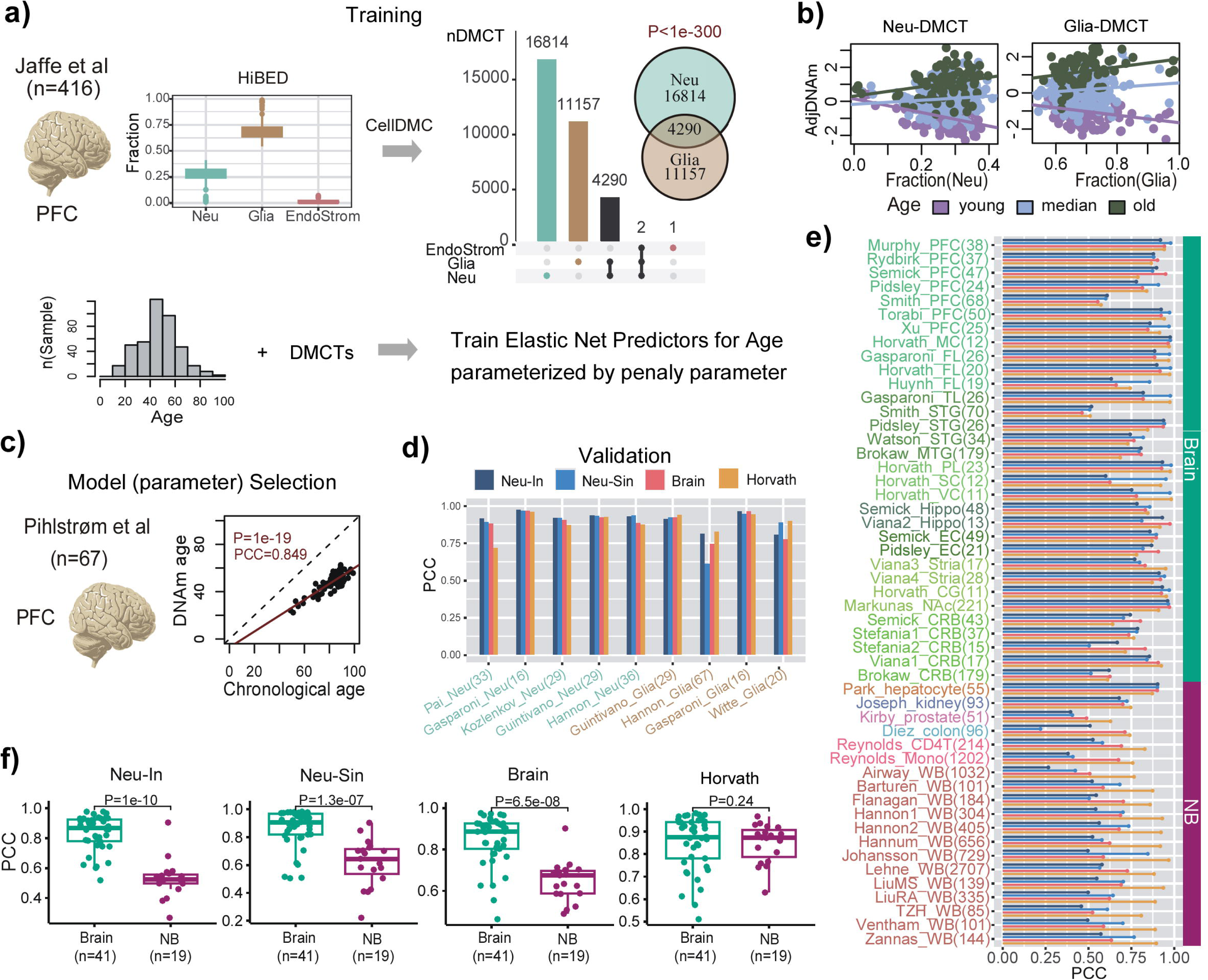
Construction and validation of neuron-clocks. **a)** The training is done on the Jaffe et al dataset encompassing 416 prefrontal cortex (PFC) samples. Fractions for 3 broad cell-types (neurons, glia and endothelial/stromal) are inferred using the HiBED algorithm. Subsequently, CellDMC is applied to infer DMCTs in each cell-type. The Fisher-test one-tailed P-value of overlap between Neu-DMCTs and Glia-DMCTs is given. Finally, elastic net predictors of chronological age are trained from the Neu-DMCTs parameterized by a penalty parameter. **b)** Scatterplots of adjusted DNAm values (adjusted for variations in cell-type fractions) of one Neu-DMCT and one Glia-DMCT against the corresponding Neuron and Glia fraction, respectively. Samples have been colored according to age-group and regression lines for each age-group have been added. **c)** Optimal model (parameter) selection is performed using a completely independent PFC dataset from Philstrom et al. Scatterplot displays the predicted vs chronological age for the optimal model. **d)** Barplot of Pearson Correlation Coefficients (PCC) for 4 clocks (Neu-In, Neu-Sin, Brain and Horvath) across sorted neuron and glia datasets. Number of samples in each dataset is given in brackets. **e)** Barplot of Pearson Correlation Coefficients (PCC) for the same 4 clocks (Neu-In, Neu-Sin, Brain and Horvath) across brain and non-brain DNAm datasets. Each cohort is labeled with the number of samples in brackets. WB=whole blood, PFC=prefrontal cortex, TL=temporal lobe, FL=frontal lobe, MTG/STG=middle/superior temporal gyrus, PL=parietal lobe, SC=sensory cortex, VC=visual cortex, MC=motor cortex, Hippo=hippocampus, EC=entorhinal cortex, Stria=striatum, CG=cingulate gyrus, NAc=nucleus accumbens, CRB=cerebellum, Mono=monocyte. **f)** For each of the 4 clocks, boxplots of PCCs comparing the values obtained in brain vs non-brain datasets. The number of datasets in each group is given below x-axis. P-value is from a one-tailed Wilcoxon rank sum test.

### Neuron and glia-specific clocks are accelerated in Alzheimer’s Disease

Next, we asked if the clocks predict age-acceleration in neurodegeneration. To this end, we collated eleven DNAm-studies of Alzheimer’s Disease encompassing two brain regions, frontal lobe (FL) and temporal lobe (TL) (**SI table S5**), amounting to a total of 213 AD cases and 185 controls in FL, and 456 cases and 341 controls in TL. As before, we first performed a meta-analysis of cell-type fractions to see if any of these fractions change in AD (**Methods**). This revealed a significant decrease of excitatory neurons and corresponding increases of oligodendrocyte/OPC and microglia fractions in AD (**SI fig.S7**). Whilst the decreased excitatory neuron fraction was observed in both TL and FL, the microglia and oligo/OPC increases were more specific to the frontal and temporal lobes, respectively (**SI fig.S8**). Thus, only the excitatory neuron fraction is observed to change with both age and AD (**Fig.2 and SI fig.S7**), although interestingly the increased fraction seen with age is reversed in AD cases. For each of the 4 clocks in each brain region we then performed a fixed and random effects meta-analysis for AD (**Methods**), revealing a weak but statistically significant age-acceleration in AD cases for the neuron-intrinsic clock in the TL, but not for brain-specific and Horvath’s clocks (**Fig.5a-b**). In the case of Horvath’s clock the association remained non-significant upon adjustment for brain cell-type fractions (**SI fig.S9**). The association of the neuron-intrinsic clock in the PFC/FL was marginally significant. Given that the I^2^ test for heterogeneity showed great consistency in the effect sizes between cohorts and regions (**Fig.5a-b**), we extended the meta-analysis to include all datasets from both regions, now revealing significant associations for both intrinsic and semi-intrinsic neuron-clocks (**Fig.5c**). Because all of the AD datasets considered in this analysis are from bulk-tissue, we wondered if a similar finding would apply to the other major cell-type in brain, i.e. glia. To this end, we used the previous Jaffe & Philstrom DNAm datasets to build and validate analogous glia intrinsic (Glia-In) and semi-intrinsic (Glia-Sin) clocks (**Methods**, **SI fig.S10, SI tables S6-S7**), subsequently computing their DNAm-Ages in the AD datasets. Interestingly, this also revealed age-acceleration effects in the glia compartment of both FL and TL regions, with the effect being strongest for the Glia-Sin clock (**Fig.5a-c**). Comparing the predicted age-acceleration of the Neu-In with the Glia-In clock, and that of the Neu-Sin with the Glia-Sin clocks, revealed moderate but significant correlations between the Neu and Glia clock estimates (**SI fig.S11**). Thus, age-acceleration patterns in neurons and glia are similar, despite the two clocks being made up of entirely distinct sets of CpGs. In summary, these data indicate that both neuron and glia specific clocks display age-acceleration in AD, and that this acceleration is more prominent in the TL glia fraction.

**Figure-5:**
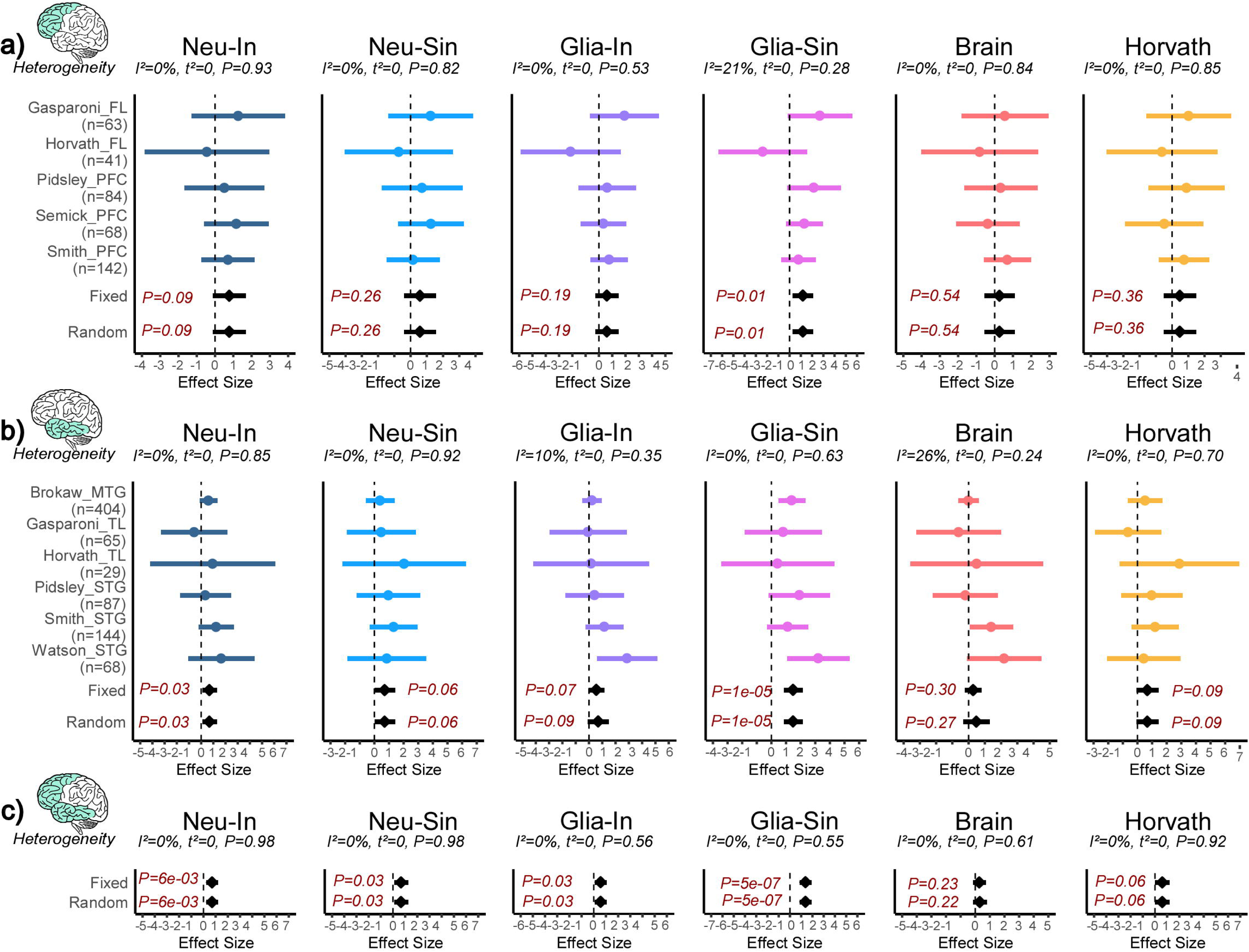
Neuron and glia specific clocks predict age-acceleration in AD. **a)** Forest plot of associations between DNAmAge and Alzheimer’s Disease (AD) for 6 different epigenetic clocks across 5 independent DNAm datasets from frontal lobe (FL) or prefrontal cortex (PFC). The effect sizes and P-values of a fixed and random effects models are also given. Statistics and P-values of heterogeneity are shown above each panel. Number of samples in each dataset is given to the left. **b)** As a) but for DNAm datasets profiling regions in the temporal lobe (TL), which includes middle and superior temporal gyrus (MTG and STG). **c)** As a), but combing FL and TL regions.

### Construction and validation of a hepatocyte-specific epigenetic clock

As a second example, we considered the case of liver tissue, aiming to build a hepatocyte-specific epigenetic clock. We used our EpiSCORE algorithm[53] and liver DNAm reference matrix defined over 5 cell-types (hepatocytes, cholangiocytes, Kupffer cells, endothelial cells and lymphocytes) [44] to estimate corresponding cell-type fractions in a series of 210 normal liver specimens (age range=18-75, mean age=47, sd=12) with available Illumina EPIC DNAm profiles [54] (**Methods**). Liver cell-type fractions changed with chronological age, although this was mainly restricted to cholangiocytes and endothelial cells, a pattern that was replicated in independent cohorts (**SI fig.S12**). Next, we assessed if we can reliably identify age-DMCTs. To this end we applied CellDMC[45] to all 210 samples, revealing that most age-DMCTs were in either hepatocytes or cholangiocytes, with hepatocyte and cholangiocyte age-DMCTs displaying strong overlap but with opposite directionality of change (**SI fig.S13a-d**). As with the age-DMCTs in neurons and glia, here too we observed a preferential co-localization to marker CpGs in our liver DNAm reference matrix (**SI fig.S5**). To validate the hepatocyte age-DMCTs, we computed the average DNAm over hepatocyte age-hypermethylated DMCTs in an independent set of 55 primary hepatocyte cultures from donors of different ages (**Methods**), which revealed the expected positive correlation with age (**SI fig.S13e**). Correspondingly, hepatocyte age-hypomethylated DMCTs displayed the expected anti-correlation, with the pattern exactly inverted for cholangiocyte age-DMTCs (**SI fig.S13e**). To further validate the hepatocyte and cholangiocyte age-DMCTs, we collected three additional liver DNAm datasets (**Methods**) and repeated the CellDMC analysis in each one. This confirmed the strong overlap between hepatocyte and cholangiocyte age-DMCTs in each validation cohort (**SI fig.S13f, SI fig.S14**). This strong validation supports the view that we can confidently identify age-associated DMCTs in hepatocytes and that these display opposite directionality of change in cholangiocytes.

Having demonstrated that we can successfully identify hepatocyte age-DMCTs, we next applied a 10-fold cross-validation procedure to build a hepatocyte-specific semi-intrinsic clock (**Methods**). Of note, this procedure involved re-applying the CellDMC algorithm on the full set of CpGs but now using only on 9 folds, with the leave-one-out bag being used for model selection (**Methods**). This procedure was iterated 10 times, each time using a different bag as the leave-one-out test set, a procedure that avoids overfitting [55], which resulted in a final age-predictor (“HepClock”) (**SI table S8, Fig.6a**). We also trained an analogous clock but now using age-DMCs derived from a linear model that only adjusts for cell-type fractions, the resulting clock thus being liver-specific but not hepatocyte-specific (“LiverClock”, **Methods, SI table S9**). To validate and benchmark these clocks, we applied them to the DNAm dataset of primary hepatocyte cultures (**Methods**). Although R-values (Pearson Correlation Coefficients) were similar for all clocks, HepClock attained a significantly lower median absolute error of 4.9 years compared to values of 20 and 10 for the liver and Horvath clocks, respectively (**Fig.6b**). This suggests that improved predictions of chronological age in sorted hepatocytes is indeed possible with a corresponding hepatocyte specific clock. Next, we applied HepClock and LiverClock to a collection of DNAm datasets from bulk-liver tissue, sorted immune-cell types, as well as other tissue-types including blood, skin, prostate, colon and kidney (**Methods**). This confirmed the clear specificity of HepClock and LiverClock to liver tissue (**Fig.6c**). We note that Horvath’s clock was excluded from this analysis, because some of the liver DNAm datasets were used for the original training of the Horvath clock itself [28].

**Figure-6:**
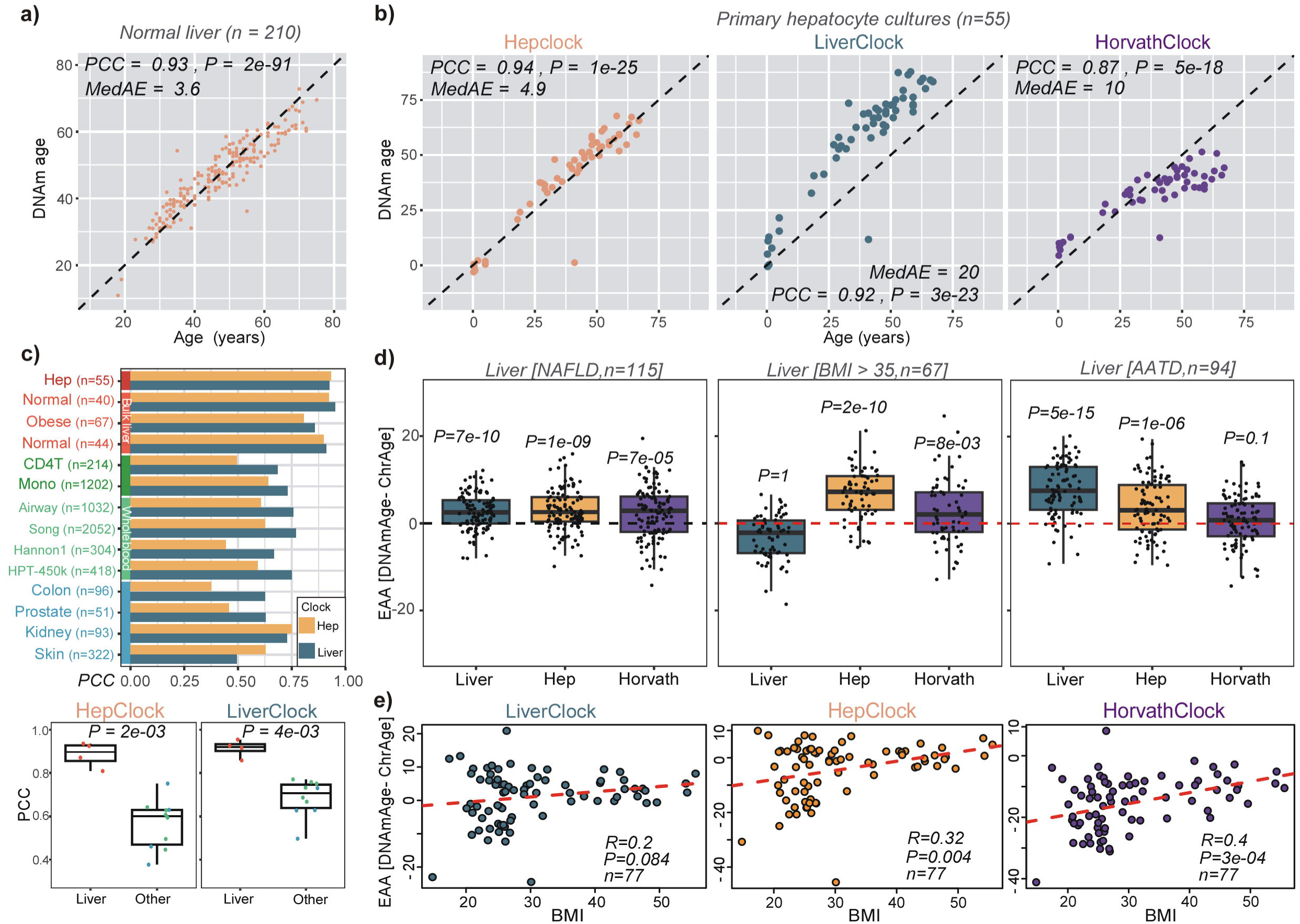
Construction and validation of hepatocyte clock. **a)** Overall strategy used to train a hepatocyte specific clock from bulk liver tissue samples. Plots show how the root mean square error (RMSE) changes as a function of the penalty parameter in the elastic net regression model, and scatterplot at the bottom compares the predicted and chronological ages for the optimal model defining the hepatocyte-clock. Pearson Correlation Coefficient (PCC), Median Absolute Error (MedAE) and regression P-value are given. **b)** Validation of the hepatocyte and liver clocks (latter trained on age-DMCs, not hepatocyte-DMCTs) in a primary hepatocyte culture. Also shown is the result for Horvath’s clock. PCC, MedAE and P-values from a linear regression are given. **c)** Barplot comparing the PCC-values of the hepatocyte and liver clocks across several liver and non-liver tissue datasets. Boxplots compare the PCC values of the hepatocyte and liver clocks in the liver/hepatocyte dataset compared to all others. P-value is from a one-tailed Wilcoxon rank sum test. **d)** Boxplots compare the extrinsic age-acceleration (EAA) of the liver, hepatocyte and Horvath clocks in two independent liver DNAm datasets, one profiling non-alcoholic fatty liver disease (NAFLD) cases and the other profiling liver-tissue from obese (BMI>35) individuals. In each case, the P-value is from a one-tailed Wilcoxon rank sum test comparing the distribution to a mean value of 0 (red dashed line). **e)** Boxplots of EAA from the three clocks in an independent DNAm liver dataset, with samples stratified according to BMI levels, as shown. For each panel, we give the PCC and linear regression test P-value.

### Hepatocyte specific clock predicts biological age-acceleration in liver pathologies

We next applied HepClock, LiverClock and Horvath’s clock to 3 separate DNAm studies representing a variety of liver pathologies, in order to assess the ability of these clocks to predict biological age-acceleration. One study encompassed 115 liver samples from patients with non-alcoholic fatty liver disease (NAFLD) [54], another set comprised 67 liver tissue samples from obese individuals (BMI>35) [56], with the 3rd cohort comprising 94 liver tissue samples from patients with Alpha-1 Antitrypsin Deficiency (AATD) [57], a genetically inherited disorder that increases the risk of cirrhosis (**Methods**). Whilst on the NAFLD samples all three clocks were accelerated, the observed acceleration was strongest for the Liver and HepClocks (**Fig.6d**). On the clinically obese samples only HepClock displayed a very significant age-acceleration effect, with Horvath’s clock being marginally associated (**Fig.6d**). In the AATD-cohort, only the Liver and HepClocks displayed age-acceleration (**Fig.6d**). In another independent liver-tissue DNAm dataset encompassing healthy non-obese and obese individuals [58, 59], HepClock and Horvath clock age-acceleration measures displayed a correlation with BMI, but not so for the LiverClock (**Fig.6e)**, mirroring the results in the previous cohort of 67 obese individuals (**Fig.6d**). The age-acceleration effect in the hepatocytes of NAFLD and obese livers is noteworthy, because the hepatocyte fraction itself decreases substantially in NAFLD and obese individuals (**SI fig.S15**). Overall, as assessed over the 4 studies, the hepatocyte-specific epigenetic clock improves the sensitivity to detect biological age acceleration in the hepatocytes of liver tissue in variety of different liver pathologies including an inherited one where Horvath’s clock did not predict age-acceleration.

### Minimal but significant overlaps with chronological age clocks

Having demonstrated how cell-type specific clocks predict age-acceleration in relevant conditions and cell-types, it is important to study the nature of the specific CpGs making up these clocks. We first assessed overlaps of our clock CpGs with previous clocks including Horvath [28], Zhang [11] and PhenoAge [3]. In general, although overlaps were small in absolute terms, some were statistically significant (**SI fig.S16**). For instance, most of our cell-type specific clock CpGs displayed a significant overlap with CpGs from Zhang and Horvath’s clocks, two clocks trained to predict chronological age. The smaller non-significant overlap with PhenoAge clock CpGs may indicate that our cell-type specific clocks in brain and liver are capturing different facets of biological aging compared to PhenoAge.

### Minimal but significant overlap with DamAge causality clock

The overlap with the recently proposed causal DamAge and AdaptAge clocks [60] was also small in absolute terms, but interestingly the neuron and glia-clock CpGs displayed a significant overlap with the DamAge clock (Binomial P < 0.001) and not with AdaptAge (Binomial P>0.05) (**SI fig.S16**). The overlapping CpGs from Neu-In and Neu-Sin with DamAge mapped to 4 genes (*SESN2, EPS8L2, SGMS1, ZNF642*), whilst the corresponding overlapping DamAge-CpGs from Glia-In and Glia-Sin mapped to 3 genes (*ST3GAL4, CBX7, TOMM40L*) (**SI table S10**). Supporting the statistical significance of these overlaps with the neuron and glia clocks, these overlapping genes have previously been implicated in AD. For instance, *SESN*2 plays an important role in AD progression [61, 62] and is regarded as a diagnostic AD biomarker [63]. The expression level of *EPS8L2* is significantly reduced in an AD mouse model [64], whilst *SGMS1* is functionally implicated in AD pathogenesis [65], displaying elevated expression levels in the brain of AD cases [66]. Interestingly, *ZNF642* is upregulated in Parkinson’s disease [67], and Zinc finger proteins are well-known to be associated with neurological diseases [68]. One of the shared DamAge glia-clock CpGs mapped to the promoter of *ST3GAL4*, a gene associated with brain atrophy in AD patients [69, 70] and that correlates with Braak staging [71]. Expression of PRC1 component *CBX7* has been shown to regulate the AD related INK4b-ARF-INK4a locus [72] and is associated with suppression of axon growth and regeneration [73]. Finally, *TOMM40L* is in linkage disequilibrium with APOE and contributes synergistically to the risk of Alzheimer’s disease [74–77]. Specifically, the *APOE*LJ*4-TOMM40’ 523 L* haplotype is associated with a higher risk and shorter times of conversion from mild cognitive impairment (MCI) to AD [78], possibly driven by mitochondrial dysfunction [75]. Overall, these data demonstrate that our brain cell-type specific clocks do map to CpGs and genes that have been causally and negatively implicated in neurodegeneration and AD, which is consistent with our neuron and glia clocks displaying age-acceleration in AD.

### The Neu-In clock CpGs are enriched for AD EWAS hits and mQTLs

Given the associations of our clocks with AD and BMI, we next asked if our clock CpGs overlap with AD or BMI associated differentially methylated cytosines (DMCs), as determined by corresponding EWAS [79, 80] (**Methods**). Whilst for the brain-clocks we did observe a significant overlap with AD-DMCs, this was not evident for the Hep/LiverClocks and BMI-DMCs (**SI fig.S17**).

Next, we asked if our clock CpGs are associated with mQTLs. We performed an enrichment against three broad classes of mQTLs: adult blood derived mQTLs as given by the ARIES database [81], mQTLs derived from the eGTEX consortium [82] which are available for a number of different tissue-types including colon, prostate, ovary, lung, breast, kidney, skeletal muscle, testis and whole blood, and mQTLs derived for fetal and adult brain [83, 84] (**Methods**). Using the ARIES database, this revealed a significant enrichment for mQTLs among brain clock CpGs, but only marginal associations for the Hep/LiverClocks (**SI fig.S18a**), probably because of the smaller number of Hep/LiverClock CpGs, itself reflecting the smaller training set in the case of liver tissue. Interestingly, all brain clock CpGs displayed significant enrichment for mQTLs as defined in adult brain, but not fetal brain, although this negative finding could be due to the smaller number of mQTLs in fetal brain (**SI fig.S18b**). For the mQTLs as defined by eGTEX we also observed enrichment, but this was restricted mainly to the neuron-clocks and tissues with the largest numbers of mQTLs (e.g. colon, lung, prostate and ovary) (**SI fig.S18c**), suggesting that other associations may become significant if we had more CpGs in either the clock or mQTL list. Supporting this, we note that we did not observe any enrichment for mQTLs as derived from whole blood in eGTEX, whilst we did observe enrichment against the larger mQTL ARIES database. In the case of the Neu-In clock and blood mQTLs from ARIES, we observed 2 clock-CpGs defining cis-mQTLs where the linked SNP has been associated with AD [85] (**SI fig.S18d, SI table S11**). Interestingly, these cis-mQTLs map to the gene *CASS4,* a CAS scaffolding protein family member involved in the formation of neuritic plaques and neurofibrillary tangles, as well as in the disruption of synaptic connections in AD [86, 87]. Of note, Neu-In clock CpGs defining cis-mQTLs in adult brain also included SNPs that mapped to *CASS4.* In addition, another Neu-In/Neu-Sin clock CpG defines a cis-mQTL in brain mapping to the *TSPAN14* gene (**SI fig.S18d**), which is associated with increased AD risk [85, 88, 89]. Thus, a minor component of the neuron-specific clocks involve CpGs whose DNAm-levels are also influenced by genetic variants associated with AD.

## Discussion

Here we have quantified, within the context of epigenetic clocks, the relative contributions of extrinsic and intrinsic aging, demonstrating that the extrinsic component in blood can account for up to around 40% of an epigenetic clock’s accuracy, whilst in brain this fraction is much lower at around 12%. Thus, whilst most of an epigenetic clock’s accuracy is intrinsic, driven by age-associated DNAm changes in individual cell-types, the age-related shifts in tissue composition can contribute a substantial amount. For instance, based on our analysis, epigenetic clocks like Horvath or Zhang clocks [11, 28], which typically attain an R^2^ value of 0.8 or higher, would likely only display R^2^ values of approximately 0.5 if one were to rederive the clocks adjusting for 12 immune-cell type fractions. Whilst in the case of blood, the extrinsic component is driven by an age-related increase of the mature to naïve T-cell fractions, in brain, this is driven by an age-related shift in the excitatory to inhibitory neuron fractions.

Our second key contribution is the dissection of the intrinsic component of aging through construction of cell-type specific epigenetic clocks. As shown here in the case of brain and liver tissue, the resulting cell-type specific clocks demonstrate clear cell-type and tissue-specificity, achieving improved predictions of chronological age in the corresponding sorted cells and tissue-types. Most importantly, our cell-type specific clocks yield improved estimates of biological age in the relevant cell-types and tissues (here brain and liver), as compared to Horvath’s multi-tissue clock and the non-cell-type specific (but tissue-specific) brain and liver clocks, the latter result highlighting the importance of distinguishing cell-type specific from tissue-specific clocks.

The case of Alzheimer’s Disease is important to discuss further, since there has been substantial controversy and debate as to whether epigenetic clocks display age-acceleration in AD or not. Here we have significantly advanced the field by constructing neuron and glia-specific clocks, and demonstrating an age-acceleration effect for both neurons and glia, with the effect being strongest for the Glia-Sin clock in the temporal lobe, suggesting that epigenetic age-acceleration associated with AD may be particularly prominent in the glia TL compartment. As far as the age-acceleration of the Neu-In and Neu-Sin clocks is concerned, it is worth noting that the excitatory neuron fraction was observed to increase with age, whilst it was decreased in AD. This pattern would suggest that the observed age-acceleration of the neuron-compartment in AD is not due to shifts in underlying neuronal subtype proportions, but instead caused by DNAm changes in these specific neuronal subtypes. It is also noteworthy that one of the CpGs in the Neu-In clock defines a cis-mQTL linked with a SNP that has been associated with AD and which maps to *CASS4*, a gene which has previously been implicated in neurotic plaque formation and loss of synaptic connectivity in AD [86, 87]. Another CpG in the Neu-In clock mapped to a cis-mQTL implicating *TSPAN14*, which is associated with increased AD risk [85, 88, 89]. These findings are important, because it is conceivable that clock-CpGs whose DNAm levels track age-acceleration in AD due to environmental risk factors (e.g. social isolation, sleep deprivation) may also be under the influence of genetic variants that determine AD-risk, which would thus point towards a shared causal pathway to AD [17].

The neuron and glia specific clock CpGs also displayed a statistically significant overlap with CpGs of the causal DamAge clocks. Remarkably, the few overlapping CpGs mapped to genes that have been strongly implicated in AD. Alongside *CBX7* and *TOMM40L* discussed earlier, it is worth highlighting here also *SESN2* and *SGMS1.* Inactivation of Sestrin-2 (*SESN2*) results in diverse metabolic pathologies, including oxidative damage, fat accumulation, mitochondrial dysfunction, and muscle degeneration, that resemble accelerated tissue aging [90, 91]. *SESN2* is also induced by amyloid-β, playing a protective role against amyloid-β neurotoxicity [61]. Knockdown of sphingomyelin synthase-1 (*SGMS1*) attenuates AD-like pathology through promoting lysosomal degradation of BACE1 [65]. Moreover, *SGMS1* is significantly elevated in the hippocampus of AD brain and inhibition significantly reduces the level of amyloid-β [66]. Of note, for *SGMS1*, the implicated CpG maps to the 5’UTR, where an age-related decrease in DNAm (as observed here in our brain clocks, **SI table S10**) would normally be associated with increased expression, hence consistent with the damaging effect as implicated in the DamAge clock. However, it should also be noted that the relation between DNAm and gene-expression is complex.

The ability to quantify biological age at cell-type resolution can also help address whether age-acceleration in various cell-types of a tissue correlate with each other or not. Here we explored this important question in the context of the neuron and glia clocks, revealing that the age-acceleration of these two clocks are indeed significantly correlated. Despite this, the age-acceleration measures were also sufficiently distinct, with the glia clocks displaying stronger acceleration in AD compared to the neuron-clocks. In the case of liver, it should be noted that most of the identified age-associated DNAm changes occurred in hepatocytes and cholangiocytes, with the underlying DNAm changes in these two cell-types displaying a very strong anti-correlation. Thus, a putative cholangiocyte clock would be composed of overwhelmingly the exact same CpGs making up the hepatocyte clock, and thus be effectively analogous to it, which is why we did not consider it here.

Whilst the computational framework presented here could be extended to construct cell-type specific epigenetic clocks for other tissue and cell-types, it is worth highlighting some of the key limitations. First, it must be stressed that our strategy hinges critically on the ability to detect age-associated DNAm changes in individual cell-types and that this requires appropriately powered datasets [26, 51]. Indeed, the need for large sample sizes (typically at least many hundreds) has recently been emphasized by us [51]. The required sample size to ensure adequate power is a complex function of many parameters, including the number of cell-types (i.e. cellular resolution) at which the inference is to be made and the distribution (notably means and variances) of their proportions across samples. At higher cellular resolution, there is a corresponding need that the cohort be big enough so that all possible combinations of cell-type fractions are realized, which in practice may be difficult because of intrinsic correlations between cell-types. Thus, the need for fairly large DNAm datasets currently limits our strategy to a few tissue-types (e.g. brain/liver) and a low cellular resolution (e.g. neuron, glia and stroma). In order to be able to quantify biological age in additional subsets, say glia subtypes in brain, or for the different immune cell subtypes in liver, one would need on the order of thousands of samples, which in future could be realized by merging DNAm datasets together. For instance, for an age-related disease like NAFLD that is associated with molecular alterations in hepatocytes and which leads to a marked reduction in the hepatocyte fraction with age, other cell-types (e.g. Kupffer macrophages) may also undergo biological aging, marking the persistent inflammation associated with NAFLD [92], but here we were underpowered to detect age-DMCTs in macrophages. Ultimately, increased cellular resolution may be critical as for instance, adjustment for a broad cell-type that encompasses two cell subtypes cannot account for age-related relative variations in these subtypes. A related important question is the relevant cellular resolution or definition of cell-types [93] to use in such analyses, as this is often dependent on the specific biological questions to be addressed.

Another key unresolved issue is the relative quantification of cell-type specific versus shared/common age-associated DNAm changes. For instance, as shown by us, most age-associated DNAm changes in blood seem to be shared between distinct immune cell-types [94], implying that the number of truly cell-type specific age-associated DNAm changes may be small in a tissue like blood. Ideally, cell-type specific clocks should be built from those CpGs that only change in one cell-type of the tissue and not others, yet confidently identifying these is challenging and would require the aforementioned very large DNAm datasets. Certainly, an in-depth study in a tissue like blood, where large numbers of DNAm datasets are available would merit further investigation, although given that the relative fractions of naïve and mature T-cell subtypes change with age [24], this may require on the order of tens of thousands of samples. In the case of brain too, we have observed a substantial overlap between neuron and glia age-associated DMCs, the resulting lineage-specificity of the corresponding clocks also being less evident than in the case of liver and the hepatocyte specific clock. Indeed, in the case of liver it is worth stressing that we were able to confidently validate the hepatocyte specific age-DMCs in sorted hepatocytes, as well as the hepatocyte and cholangiocyte age-DMCs in 3 independent bulk liver DNAm datasets.

In summary, we have quantified the relative contributions of extrinsic and intrinsic aging to an epigenetic clock’s predictive accuracy, whilst also presenting a computational strategy to dissect the intrinsic component, building cell-type specific epigenetic clocks that clearly demonstrate cell-type and tissue specificity in relation to prediction of chronological and biological age. The construction of such cell-type specific clocks will be critical to advance our understanding of biological aging at cell-type resolution and for evaluating in which cell-types cellular rejuvenation interventions work best.

## Methods

### Brief description of main datasets analyzed

#### Human liver DNA methylation datasets

##### Johnson et al [54]

This is an Illumina EPIC DNAm dataset of liver-tissue, comprising 341 samples, with 16 duplicates among them. The dataset includes samples from four stages representing the progression of non-alcoholic fatty liver disease (NAFLD): stage 0, stage 3, stage 3-4, and stage 4. Stage 0 is normal liver. Data is available at GEO (http://www.ncbi.nlm.nih.gov/geo/; accession number GSE180474). We downloade a preprocessed data matrix following QC, including P-values of detection for each probe. To select high-quality probes for clock construction, we utilized the dropLociWithSnps() function of minfi v.1.44.0 to remove SNP probes [95]. Sex chromosome CpGs were also excluded to minimize gender disparities. Any beta value with a P-value greater than 0.01 was considered as a missing value, and any probe with a coverage-fraction less than 0.99 was discarded. Missing values were then imputed using impute.knn (k=5) of impute v.1.72.3 [96]. Type-2 bias was corrected using BMIQ algorithm [97], and finally duplicates were averaged, resulting in a final data matrix of 325 samples encompassing 812,168 CpGs on autosomes. Of these 325 samples, 210 were from normal healthy liver, whereas the remaining 115 represent NAFLD of grade-3 or higher. In this work, the 210 normal liver samples were used to build and validate the Liver and HepClocks in a 10-fold CV-strategy, whereas the 115 NAFLD samples were used to test for biological age-acceleration.

##### Park et al [98]

Illumina HM850 (EPIC) dataset encompassing 56 primary hepatocyte cultures. Raw IDAT files were downloaded from GSE123995. Data was processed as before. Any probe with missing values was excluded, resulting in 809,289 probes. A sample was removed due to very low predicted hepatocyte fraction (as estimated using EpiSCORE).

##### Bonder et al [56]

Illumina HM450k dataset profiling 67 obese liver samples. All individuals have a BMI greater than 35. Raw IDAT files were downloaded from GSE61446. minfi::preprocessRaw() was employed to process the raw IDAT data. The remaining steps in the preprocessing pipeline align with the aforementioned description, with a probe coverage threshold set at 0.90. A total of 411,233 probes remained.

##### Bacalini et al [99]

Illumina HM450 dataset profiling 40 normal liver samples. Raw IDAT files were downloaded from GSE107038 and processed as the other datasets above. Any probe with missing values was excluded, resulting in 446,651 probes. A sample was removed due to very low predicted hepatocyte fraction (as estimated using EpiSCORE).

##### Horvath and Ahrens et al [58, 59]

Illumina HM450 datasets comprising 77 samples from healthy non-obese (n=44) and healthy obese (n=33) individuals. Processed data was downloaded from GEO under accession numbers GSE61258 and GSE48325. For both datasets, we only used probes with no missing values (i.e. significant detection P-value < 0.01) across all samples. Datasets were then normalized with BMIQ. The merged dataset encompassed 77 samples and 417,213 probes. A subset of samples were obtained following bariatric surgery, but since surgery had no effect on clock estimates [58, 59] we kept these samples in. Of note, the subset of 44 healthy non-obese samples were used to assess the predictive accuracy of the clocks with chronological age.

##### Wang et al AATD dataset [57]

Illumina HM850 (EPIC) dataset encompassing 108 liver samples from 94 Alpha-1 Antitrypsin Deficiency (AATD) patients and 14 AATD patients with cirrhosis. Only the normal liver samples from the 94 AATD patients without cirrhosis were used. Processed normalized data was downloaded from GEO under accession number GSE119100. This normalized data is already adjusted for type-2 probe bias (adjusted with Subset-quantile Within Array Normalization (SWAN)), and probes that failed to achieve significant detection (p<0.01) in all samples alongside cross-reactive probes were filtered out, leaving a total of 866,836 probes.

#### Human brain DNA methylation datasets used for training and model selection

##### Jaffe

The 450k dataset from Jaffe et al [50], which profiled DNAm in human dorsolateral prefrontal cortex from controls and patients with schizophrenia (SZ), was obtained from the NCBI GEO website under the accession number GSE74193 (https://www.ncbi.nlm.nih.gov/geo/query/acc.cgi?acc=GSE74193). The file “GSE169156_RAW.tar” which contains the IDAT files was downloaded and processed with minfi package [95]. Probes with >25% NAs (defined by PC>C0.01) were discarded. Samples younger than 18 years old were also discarded. The filtered data was subsequently normalized with BMIQ [97], resulting in a normalized data matrix for 473,536 probes and 416 samples, of which 226 are controls and 190 are SZ cases, age range 18-97 years.

##### Pihlstrøm

The EPIC dataset from Pihlstrøm et al [52], which profiled DNAm in human frontal cortex, was obtained from the NCBI GEO website under the accession number GSE203332 (https://www.ncbi.nlm.nih.gov/geo/query/acc.cgi?acc=GSE203332). The file “GSE203332_RAW.tar” which contains the IDAT files was downloaded and processed with minfi package. Probes with NAs (defined by PC>C0.01) were discarded. The filtered data was subsequently normalized with BMIQ, resulting in a normalized data matrix for 832,899 probes and 442 samples, of which 67 are controls with an age-range of 49 to 99 years.

#### Human brain DNA methylation datasets used for validation

Due to the large number of brain DNAm datasets analyzed, these are all summarized in **Supplementary Methods**.

#### Human DNA methylation datasets from other tissues

All the whole blood datasets used here have been preprocessed and analyzed previously by us [24]. In addition to whole blood, we also analyzed the following DNAm datasets from other tissue-types:

##### Colon [100]

Illumina HM450 dataset comprising 96 normal colon mucosa. Raw data is available from GSE131013. IDAT files were downloaded and processed with minfi package. Probes with NAs (defined by P > 0.01) were discarded. The filtered data was subsequently normalized with BMIQ, resulting in a normalized data matrix for 448,286 probes and 240 samples. 144 tumor samples were discarded.

##### Prostate [101]

Illumina HM450 dataset comprising 51 normal prostate samples. Raw data is available from GSE76938. IDAT files were downloaded and processed with minfi package. Probes with NAs (defined by P > 0.01) were discarded. The filtered data was subsequently normalized with BMIQ, resulting in a normalized data matrix for 471,112 probes and 123 samples and 72 prostate cancer samples were discarded.

##### Kidney [102]

Illumina HM450 dataset profiling 31 normal kidney tissue samples. Raw data is available from GSE79100. There are three technical replicates per tissue. IDAT files were downloaded and processed with minfi package. Probes with NAs (defined by P > 0.01) were discarded. The filtered data was subsequently normalized with BMIQ, resulting in a normalized data matrix for 454,671 probes and 93 samples. Skin [103]: Illumina HM450 dataset profiling 322 normal skin samples from a peri-umbilical punch biopsy. From GSE90124, we downloaded the processed type-2 probe bias adjusted DNAm data with 450,531 probes and 322 samples.

### Estimating cell-type fractions in whole blood, liver and brain

For a given bulk-tissue sample with a genome-wide DNAm profile, it is possible to estimate the cell-type composition of the tissue with a corresponding tissue-specific DNAm reference matrix [104]. Such a DNAm reference matrix consists of cell-type specific CpGs and cell-types, and is typically built from genome-wide DNAm data of sorted cell-types [105]. The cell-type fractions themselves are estimated by regressing the DNAm profile of the bulk-tissue samples against the reference profiles of the DNAm reference matrix, a procedure that only uses the highly discriminatory cell-type specific CpGs as defined in the DNAm reference matrix [106]. For whole blood, we estimated fractions for 12 immune cell subtypes (naïve and memory B-cells, naïve and memory CD4+ T-cells, naïve and memory CD8+ T-cells, T-regulatory cells, natural killer cells, monocytes, neutrophils, eosinophils, basophils) using EpiDISH and our validated DNAm reference matrix for these 12 cell-types [24]. In the case of liver tissue we used our validated EpiSCORE algorithm and the associated liver DNAm reference matrix defined for hepatocytes, cholangiocytes, endothelial cells, Kupffer macrophages and lymphocytes [53]. In the case of brain-tissue, we applied the HiBED algorithm [46], since the EpiSCORE brain DNAm reference matrix led to the microglia fraction always being zero. We applied HiBED to estimate proportions for 7 brain cell-types including GABAergic excitatory neurons, glutamatergic inhibitory neurons, astrocytes, microglia, oligodendrocytes, endothelial cells and stromal cells. When applying CellDMC we collapsed these estimates to 3 broad cell-types: neurons, glia and endothelial/stromal.

### Relative quantification of extrinsic and intrinsic aging in whole blood and brain

#### Whole Blood

As training data we used two of the largest whole blood DNAm datasets, the Airway [107] and Lehne [108] cohorts, which we have previously normalized and analyzed [24]. Briefly, the Airway EPIC DNAm dataset (GSE147740) included 1032 samples with age information. The Lehne dataset is a 450k DNAm dataset (GSE55763) encompassing 2707 samples in total. The two datasets were merged, resulting in 314,436 CpGs and 3739 samples. In the Airway dataset, age is cofounded by gender (Pearson correlation test, P-value =0.003896). To train the unadjusted and adjusted clocks, we applied the Elastic Net regression to the training dataset with alpha=0.5 and a 10-fold cross-validation to infer the optimal penalty parameter lambda. In the unadjusted case, we applied the Elastic Net to the standardized DNAm residuals after adjusting for sex and cohort only. In the adjusted case, we applied the Elastic Net to the standardized DNAm residuals after adjusting for sex, cohort and the 12 estimated immune-cell type fractions. We also trained another clock adjusted for 9 immune cell-type fractions by merging the naïve and memory lymphocyte subsets together, in other words, by adding the fractions of naïve and memory B-cells to yield a fraction for B-cells, and similarly for CD4T and CD8T-cells. The 3 optimal clocks were then applied to 11 independent whole blood datasets (using only control/healthy samples, see **SI table.S1**), all previously normalized as in Luo et al [24]. In the case of the adjusted clocks, the clocks were applied to the standardized DNAm residuals after adjustment for the 12 or 9 immune-cell fractions. Finally, the predicted ages were linearly regressed to the true chronological ages in each cohort, resulting in R^2^ values for the unadjusted and the two adjusted clocks. The ratio of the adjusted clock’s R^2^ value to that of the unadjusted clock, 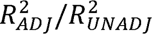 quantifies the fraction of the unadjusted clock’s accuracy that can be attributed to intrinsic aging. For the extrinsic component, its fractional contribution is given by 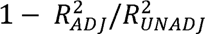.

#### Brain

For training, we used the Jaffe et al dataset [50] encompassing 491 samples, including 300 controls and 191 samples from schizophrenia patients, with an age range 0.003 ∼ 96.98 years. We used HiBED to estimate the fractions for 7-cell-types (ExN, InN, Oligo/OPC, Micro, Astro, Endo, Strom), as described earlier. To train the Elastic Net clocks with alpha=0.5, we used a 10-fold cross-validation to infer the optimal penalty parameter lambda. In the unadjusted case, we applied the Elastic Net to the standardized DNAm residuals obtained after adjusting for sex and disease-status, whilst in the adjusted case, it was applied on the standardized DNAm residuals obtained after adjusting for sex, disease-status and 7 brain cell-type fractions. The two clocks were then applied to 11 independent brain DNAm datasets (using control/healthy samples only, see **SI table.S1**), obtaining in each cohort corresponding 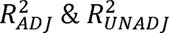 values. The fraction of the unadjusted clock’s accuracy attributed to intrinsic and extrinsic aging was then estimated as described above for whole blood.

### Construction of the neuron/glia/brain-specific epigenetic clocks

We used HiBED [46] to estimate fractions for 7 brain cell-types in the Jaffe and Philstrom datasets, in each case summing up fractions to yield estimates for 3 broad cell-types: neurons (Neu), glia (Glia) and endothelial/stromal. We then applied CellDMC [45], as implemented in the EpiDISH package [109], to the Jaffe dataset in order to infer age-associated DMCTs in each of these 3 broad cell-types. Briefly, the CellDMC model postulates the following relationship for the DNAm level 1_cs_ of a CpG *c* in sample *s*

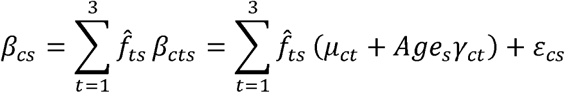

where *f̂_ts_* are the estimated cell-type fractions for cell-type *t* in sample *s*. This can be re-expressed as a multivariate regression model with interaction terms between age and cell-type fractions

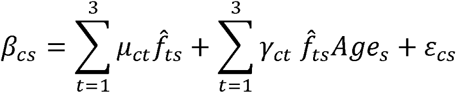

This estimates the regression coefficients *µ_ct_* and *γ_ct_* so as to minimize the error in a least squares sense. When applying CellDMC to the Jaffe dataset, we also adjusted for schizophrenia (SZ) status and gender, i.e we ran the model:

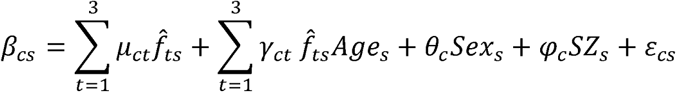

We included both SZ cases and controls, adjusting for case/control status, so as to increase power. Having identified the significant neuron-DMCTs (those CpGs *c* with a significant *γ_ct_* with *t=neuron*, t-test FDR<0.05), we next trained Elastic Net regression (glmnet R package) [49] models (alpha=0.5) for age, each parameterized by a different penalty parameter (lambda) value. This training was done on the residuals obtained by regressing out the cell-type fractions from the DNAm beta matrix. To select the optimal model, we then applied the age-predictors for each choice of lambda to the Philstrom DNAm dataset (using the residuals after adjusting this DNAm data matrix for cell-type fractions), selecting the lambda that minimized the root-mean-square error (RMSE). This defines a neuron-specific epigenetic clock that is applicable to residuals after adjusting for cell-type fraction, a clock we call “Neu-In” (neuron-intrinsic).

We also constructed a similar clock called “Neu-Sin” (neuron semi-intrinsic), following the same procedure as before, but now not adjusting the DNAm data matrix for cell-type fractions. In other words, whilst the Neu-In and Neu-Sin clocks are both built from CpGs that change with age in neurons, the Neu-In clock uses the residuals for construction, whilst the Neu-Sin clock does not. A third non-cell-type specific clock (Brain) was also built using the same procedure as before, but not using CellDMC to identify CpGs changing with age, instead identifying age-DMCs by running a linear model of DNAm versus age adjusting for cell-type fractions, i.e. in this case the regression model is

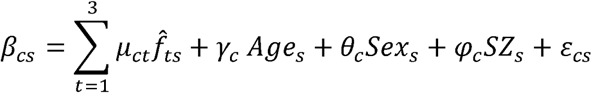

and one finds age-DMCs, as those CpGs *c* with a significant *γ_c_* with *t=hepatocyte*, (t-test, FDR<0.05). The age-predictor itself was then also trained on residuals obtained after regressing out the effect of cell-type fractions. Of note, the model selection step to define the Neu-Sin and Brain clocks was also done using the independent Philstrom DNAm dataset.

Finally, an analogous procedure describing the construction of the Neu-In and Neu-Sin clocks was followed to build corresponding glia-specific intrinsic and semi-intrinsic clocks (Glia-In & Glia-Sin).

### Validation of the neuron/glia/brain specific clocks

To validate and demonstrate specificity of the Neu-In/Sin, Glia-In/Sin and Brain clocks, we applied them to sorted neuron, sorted glia, brain and non-brain DNAm cohorts. When applying the Neu-In, Glia-In and Brain clocks, we adjusted the DNAm datasets for underlying cell-type fractions, which were estimated using HiBED in the case of brain tissue, EpiDISH in whole blood cohorts and EpiSCORE in other non-brain tissues.

### Construction of HepClock and LiverClock

We first describe the construction of the hepatocyte-specific clock (HepClock). We selected 210 normal liver samples from Johnson et al.’s dataset for training and model selection. EpiSCORE was used to estimate fractions for 5 liver cell-types including hepatocytes, as described above. The 210 samples were then randomly divided into 10 bags (each bag with 21 samples), with stratified sampling within each age group to ensure a similar age distribution in each bag. To identify CpGs changing with age in hepatocytes, we then applied the CellDMC algorithm [45] to a subset composed of 9 bags, resulting in a set of hepatocyte age-DMCTs. The specific CellDMC model run was

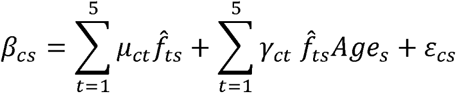

Having identified the significant hepatocyte-DMCTs (those CpGs *c* with a significant r_ct_ with *t=hepatocyte,* t-test FDR<0.05), and using the same 9 bags, we then trained a range of lasso regression models parameterized by a penalty parameter lambda. Each of these models (parameterized by lambda) were subsequently applied to the left-out-bag to yield age-predictions for the samples in this left-out-bag. This procedure was repeated 10-times, each time using a different subset of 9 bags for training and one left-out-bag for model selection. The age-predictions over all 10 folds were then combined and compared to the true ages computing the RMSE for each choice of lambda. The model minimizing the RMSE was then selected as the optimal model. Of note, the above procedure applies CellDMC a total of 10 times, once for each fold. The number of hepatocyte age-DMCTs is variable between folds, so we identified the minimum number (k) obtained across all folds, and subsequently only used the top-k hepatocyte age-DMCTs from each fold before running the lasso regression model. Lasso was implemented using the glmnet function from the glmnet R-package (version 4.7.1) [49], with alpha set to 1. Having identified the optimal model (lambda parameter), we finally reran the lasso model with this optimal parameter on the full set of 210 samples using the top-k hepatocyte age-DMCTs as inferred using CellDMC on all samples.

The LiverClock was built in an analogous fashion with only one key difference: instead of applying CellDMC in each fold, we ran a linear regression of DNAm against age with all the liver cell-type fractions as covariates. That is, the model in this case is

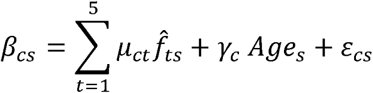

and one finds age-DMCs, as those CpGs *c* with a significant *γ_c_* (t-test, FDR<0.05). After optimization of lambda, the final age lasso predictor was built from such age-DMCs derived using all 210 samples.

### Simulation analysis on mixtures of sorted cell-types

We used a simulation framework to assess and compare the intrinsic and semi-intrinsic paradigms for cell-type specific clock construction. We first generated basis DNAm data matrices for 3 hypothetical cell-types “A”, “B” and “C” each one defined over 10,000 CpGs and 200 samples, drawing DNAm beta-values from Beta(10,40), i.e. from a Beta-distribution with a mean value of 0.25. We then declared the first 1000 CpGs to be cell-type specific for a cell-type “A”, by drawing the DNAm-values for two other cell-types (“B” and “C”) from Beta(40,10), i.e. from a Beta-distribution with mean 0.75. Thus, the first 1000 CpGs are unmethylated in cell-type “A” compared to “B” and “C”. Likewise, another 1000 non-overlapping CpGs were defined to be unmethylated in cell-type “B” and another 1000 non-overlapping CpGs for cell-type “C”. The remaining 7000 CpGs are unmethylated in all cell-types. To generate realistic mixtures of these 3 cell-types, we randomly sampled cell-type fractions from true estimated fractions in whole blood by applying our EpiDISH algorithm [109] to estimate 7 immune cell-type fractions in a large dataset of 656 whole blood samples [110]. We declared 3 cell-types to be granulocytes, monocytes and lymphocytes, by adding together the fractions of B-cells, CD4T-cells, CD8 T-cells and NK-cells to represent “lymphocytes” and adding together the fractions of neutrophils and eosinophils to represent “granulocytes”. To define the ages of the 200 samples we bootstrapped 200 values from the ages of 139 donors from the BLUEPRINT consortium [111] in order to mimic an age distribution of a real human cohort. Before mixing together the artificially generated DNAm data matrix for the 3 hypothetical cell-types, we introduced age-DMCTs in the cell-type defining lymphocytes. This was done by drawing DNAm-values for these age-DMCTs to increase with the age of the sample. In more detail, because the minimum age of the BLUERPINT samples was 30, we first assigned M-values according to the formula:

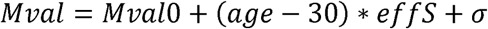

where *effS=0.1* and *Mval0=log2(0.1/(1-0.1))* and a representing a Gaussian deviate with a standard deviation value equal to *effS*. These values were chosen to mimic age-related DNAm changes as seen in real data, although with a smaller noise component in order to better understand and interpret downstream results.

In total we considered 4 different scenarios depending on the choice of age-DMCTs in lymphocytes and whether the fraction of “lymphocytes” increases with age or not. In scenarios (i) and (ii), 1000 age-DMCTs were chosen to not overlap with any of the 3000 cell-type specific marker CpGs. In scenarios (iii) and (iv), 900 age-DMCTs in lymphocytes were chosen to be lymphocyte-specific markers. Reason we did not pick all 1000 lymphocyte markers, is to leave 100 markers that remain cell-type specific to allow for reference construction and estimation of cell-type fractions using EpiDISH. In scenarios (i) and (iii), cell-type fractions were chosen so that no fraction changed significantly with age, whilst in scenarios (ii) and (iv) the lymphocyte-fraction was chosen to increase with age.

For each of the 4 scenarios, we generated training and validation mixture sets. DNAm reference matrices were built from different realizations of the basis DNAm data matrices but using the same parameter values. These reference matrices were then applied to the training dataset to infer cell-type fractions. With the estimated cell-type fractions we then applied CellDMC [45] to the training set to infer age-DMCTs. Since age-DMCTs are only introduced in lymphocytes, we focus on the predicted age-DMCTs in this cell-type, applying an Elastic Net (alpha=0.5) penalized regression framework in a 10-fold cross-validation framework to learn a predictor of chronological age using either an intrinsic or semi-intrinsic paradigm: in the semi-intrinsic paradigm the Elastic Net predictor was trained on the original DNAm-values, whilst in the intrinsic case it was applied to residuals obtained by regressing out the effect of the cell-type fractions. The optimal penalty parameter was selected via the 10-fold CV procedure and the optimal predictor then applied to the validation dataset. Of note, in the semi-intrinsic setting the validation set is defined on the original DNAm-values, whereas the intrinsic model is applied to the residuals of the validation set. For each scenario we ran 50 different simulations, recording the accuracy (root mean square error-RMSE and Pearson Corrrelation Coefficient-PCC) of the semi-intrinsic and intrinsic lymphocyte-specific clock predictors in the corresponding validation set.

### Meta-analyses

In the case of brain tissue datasets, we performed two separate meta-analyses. One of these explored if brain cell-type fractions change with age, whilst the other was done to explore if our cell type specific clocks (extrinsic age-acceleration-EAA) display correlations with Alzheimer’s Disease (AD) status. The former analysis was done for 7 brain cell types over 13 independent DNAm datasets encompassing frontal and temporal lobe regions, in each dataset adjusting for disease status and sex. The latter meta-analysis was done for frontal lobe (n=5) and temporal lobe (n=6) separately, and then again treating both regions together (n=11). In each cohort, associations between EAA and AD status was assessed using a multivariate linear regression with sex and age as covariates, AD status encoded as ordinal with 0=control, 1=AD case. We extracted the corresponding effect size, standard error, Student’s t test, and P-value for each regression, Meta-analysis was then performed using both fixed effect and random effect inverse variance method using the metagen function implemented in the meta R-package [112, 113].

### Co-localization of age-DMCTs with DNAm reference matrix CpGs

The liver DNAm reference matrix used to estimate cell type proportions before training the clocks contains 171 marker gene promoters. We calculated the number of hepatocyte and cholangiocyte age-DMCTs that mapped to the promoter regions of these 171 marker genes (where the region is defined as within 1kb upstream or downstream of the transcription start site). To generate a null distribution we did the following: from the DNAm dataset that we used to infer the age-DMCTs, we randomly selected a matched number of age-DMCTs and checked how many mapped to the promoter regions of these 171 marker genes. This process was repeated 1,000 times to obtain the null distribution, yielding an empirical P-value. In the case of the HiBED DNAm reference matrix, we focused on the CpGs in the first layer of deconvolution, and computed overlaps with neuron and glia age-DMCTs using a 1kb window. A null distribution was estimated in the same way as for liver.

### Enrichment analyses

We assessed enrichment of clock-CpGs against EWAS-DMCs for AD and BMI, as determined by the EWAS catalog [79] and EWAS-atlas [80]. Enrichment analysis was done using a one-tailed Fisher exact test. We also assessed enrichment of our clock-CpGs for (i) blood mQTLs as determined by the ARIES mQTL database (n=123010 is the number of distinct CpGs in mQTLs present in the training cohorts of the clocks) [81], focusing on the mQTLs identified in the adult population, (ii) mQTLs (n[breast]=13256, n[colon]=156419, n[kidney]=18578, n[lung]=166242, n[skeleton muscle]=14437, n[ovary]=134215, n[prostate]=68724, n[testis]=14894, n[blood]=21222) identified by the eGTEX consortium [82] and (iii) mQTLs identified from adult [84] (n= 71965) and fetal [83] (n= 30707) brain DNAm datasets. Numbers of mQTLs mapping to the datasets used to train the clocks was computed. Since the GTEx project identified mQTLs using the Illumina EPIC array, while our clocks were trained using 450k probes, we only used mQTLs that map to 450k probes. Significance was evaluated using a one-tailed Binomial test. Overlap with GWAS AD SNPs from Bellenguez et al [85] and GWAS BMI SNPs from Pulit et al [114] was done by restricting to SNPs that were part of mQTLs that included clock-CpGs.

## Code Availability

All the cell-type specific clocks presented here are freely available as part of the R-package CTSclocks, downloadable from https://github.com/HGT-UwU/CTSclocks

## Data Availability

The Illumina DNA methylation datasets analyzed here are all freely available from GEO (www.ncbi.nlm.nih.gov/geo). Details: Liver-NAFLD (n=325, GEO: GSE180474); Hepatocyte cultures (n=56, GEO: GSE123995); Liver-obese (n=67, GEO: GSE61446); Liver-Horvath/Ahrens (n=77, GEO: GSE61258 & GSE48325); Liver tissue (40 normal, GEO: GSE107038); Liver-AATD (n=94, GEO: GSE119100); MESA (214 purified CD4+ T-cells and 1202 Monocyte samples, GEO: GSE56046 & GSE56581); Colon (96 normal and 144 adenocarcinoma, GEO: GSE131013); Prostate (123 normal and 63 benign, GEO: GSE76938); Kidney (93 normal, GEO: GSE79100); Skin (322 normal, GEO: GSE90124); Brain sorted-Pai (33 normal and 67 BD, SCZ, GEO: GSE112179); Brain sorted-Gasparoni (32 normal and 30 AD, GEO: GSE66351); Brain sorted-Kozlenkov (29 normal and 59 Heroin or suicide, GEO: GSE98203); Brain sorted-Guintivano (58 normal, GEO: GSE41826); Brain sorted-Hannon (103 normal, GEO: GSE234520); Brain sorted-Witte (20 normal and 36 BD, SCZ, MDD, GEO: GSE191200); Brain-Jaffe (226 normal and 190 SCZ, GEO: GSE74193); Brain-Philstrom (67 normal and 375 Various mental disease and neurodegenerative disease, GEO: GSE203332); Brain-Semick (187 normal and 82 AD, GEO: GSE125895); Brain-Pidsley (71 normal and 179 AD, GEO: GSE43414); Brain-Smith (138 normal and 148 AD, GEO: GSE80970); Brain-Horvath (95 normal and 205 AD, HD, GEO: GSE72778); Brain-Gasparoni (52 normal and 76 AD, GEO: GSE66351); Brain-Watson (34 normal and 34 AD, GEO: GSE76105); Brain-Brokaw (358 normal and 450 AD, GEO: GSE134379); Brain-Murphy (38 normal and 37 MDD, GEO: GSE88890); Brain-Rydbirk (37 normal and 41 MSA, GEO: GSE143157); Brain-Torabi (50 normal and 75 SCZ, GEO: GSE128601); Brain-Xu (25 normal and 23 AUD, GEO: GSE49393); Brain-Huynh (19 normal and 27 Multiple sclerosis, GEO: GSE40360); Brain-Viana2 (13 normal and 14 SCZ, GEO: GSE89703); Brain-Viana3 (17 normal and 16 SCZ, GEO: GSE89705); Brain-Viana4 (28 normal and 21 SCZ, GEO: GSE89706); Brain-Markunas (221 normal, GEO: GSE147040); Brain-Viana1 (17 normal and 16 SCZ, GEO: GSE89702); Brain-Stefania1 (37 normal and 21 Suicide, GEO: GSE137222); Brain-Stefania2 (15 normal and 18 Suicide, GEO: GSE137223); Whole blood-Airway (1032 normal, GEO: GSE147740); Whole blood-Barturen (101 normal and 473 SARS, GEO: GSE179325); Whole blood-Flanagan (184 normal, GEO: GSE61151); Whole blood-Hannon1 (304 normal and 332 SCZ, GEO: GSE80417); Whole blood-Hannon2 (405 normal and 260 SCZ, GEO: GSE84727); Whole blood-Hannum (656 normal, GEO: GSE40279); Whole blood-Johansson (729 normal, GEO: GSE87571); Whole blood-Lehne (2707 normal, GEO: GSE55763); Whole blood-LiuMS (139 normal and 140 Multiple Scleroris, GEO: GSE106648); Whole blood-LiuRA (335 normal and 354 Rheumatoid arthritis, GEO: GSE42861); Whole blood-TZH (85 normal and 620 Different health conditions, GEO: Requested Access); Whole blood-Ventham (101 normal and 279 Inflammatory bowel disease, GEO: GSE87648); Whole blood-Zannas (144 normal and 278 Trauma, GEO: GSE72680). The DNAm-atlas encompassing DNAm reference matrices for 13 tissue-types encompassing over 40 cell-types is freely available from the EpiSCORE R-package https://github.com/aet21/EpiSCORE

## Declarations

### Ethics approval and consent to participate

Not applicable here since we have only analyzed existing publicly available data.

### Authors contributions

TH, GX and AET performed the statistical and bioinformatic analyses. AET conceived the study, and wrote the manuscript with contributions from TH and GX. QL contributed to the data-analyses. MJ and NE contributed pointers to data and helped revise the manuscript.

## Acknowledgements

We thank everyone who supports open-access data.

