## Supplementary Figures for "Cell-type specific epigenetic clocks to quantify biological age at cell-type resolution"

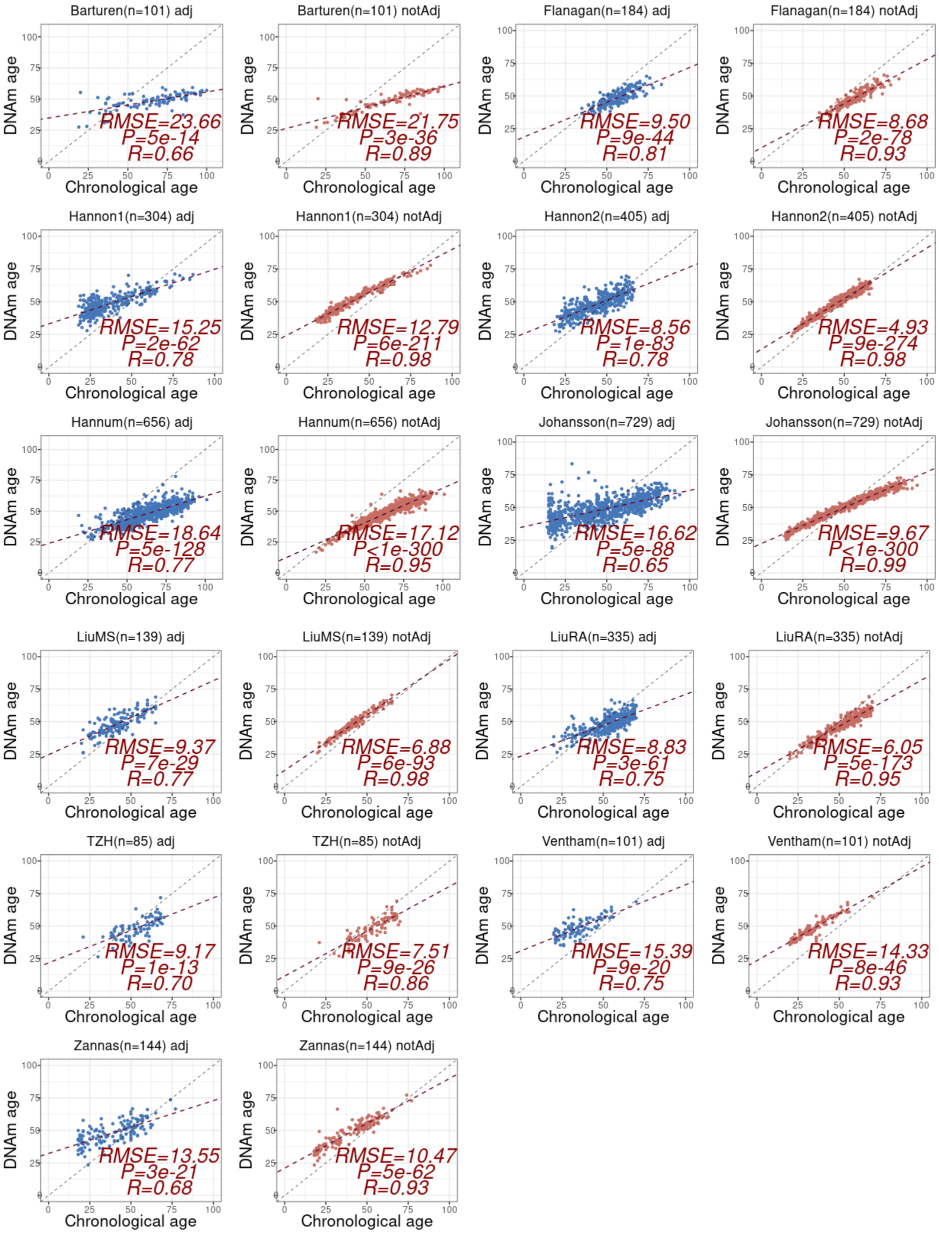


**SI fig.S1: Predicted DNAmAge vs chronological age for CTH adjusted (12 immune-cell types) and unadjusted clocks in 11 whole blood cohorts.** Each scatterplot is labeled by cohort, cohort-size and whether it is adjusted or unadjusted clock. Root Mean Square Error (RMSE), two-tailed P-value of a linear regression and R (Pearson Correlation Coefficient) value is given. Adjusted clock was adjusted for variations in naïve + memory B-cells, naïve + memory CD4T, naïve + memory CD8T-cells, T-regulatory cells, NK-cells, monocytes, neutrophils, eosinophils and basophils.

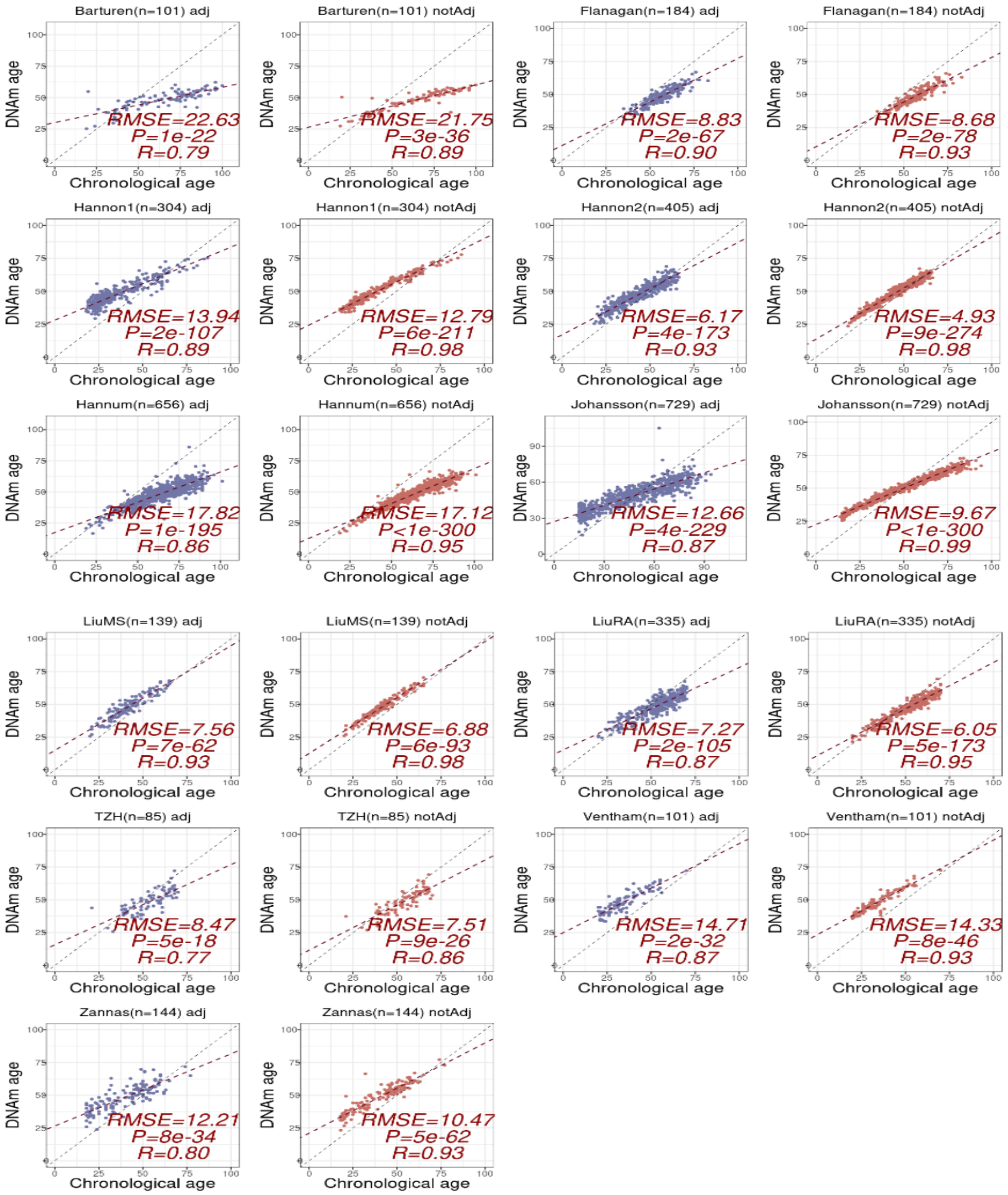


**SI fig.S2: Predicted DNAmAge vs chronological age for CTH adjusted (9 immune-cell types) and unadjusted clocks in 11 whole blood cohorts.** Each scatterplot is labeled by cohort, cohort-size and whether it is adjusted or unadjusted clock. Root Mean Square Error (RMSE), two-tailed P-value of a linear regression and R (Pearson Correlation Coefficient) value is given. Adjusted clock was adjusted for variations in B-cells, CD4T and CD8T-cells, T-regulatory cells, NK-cells, monocytes, neutrophils, eosinophils and basophils.

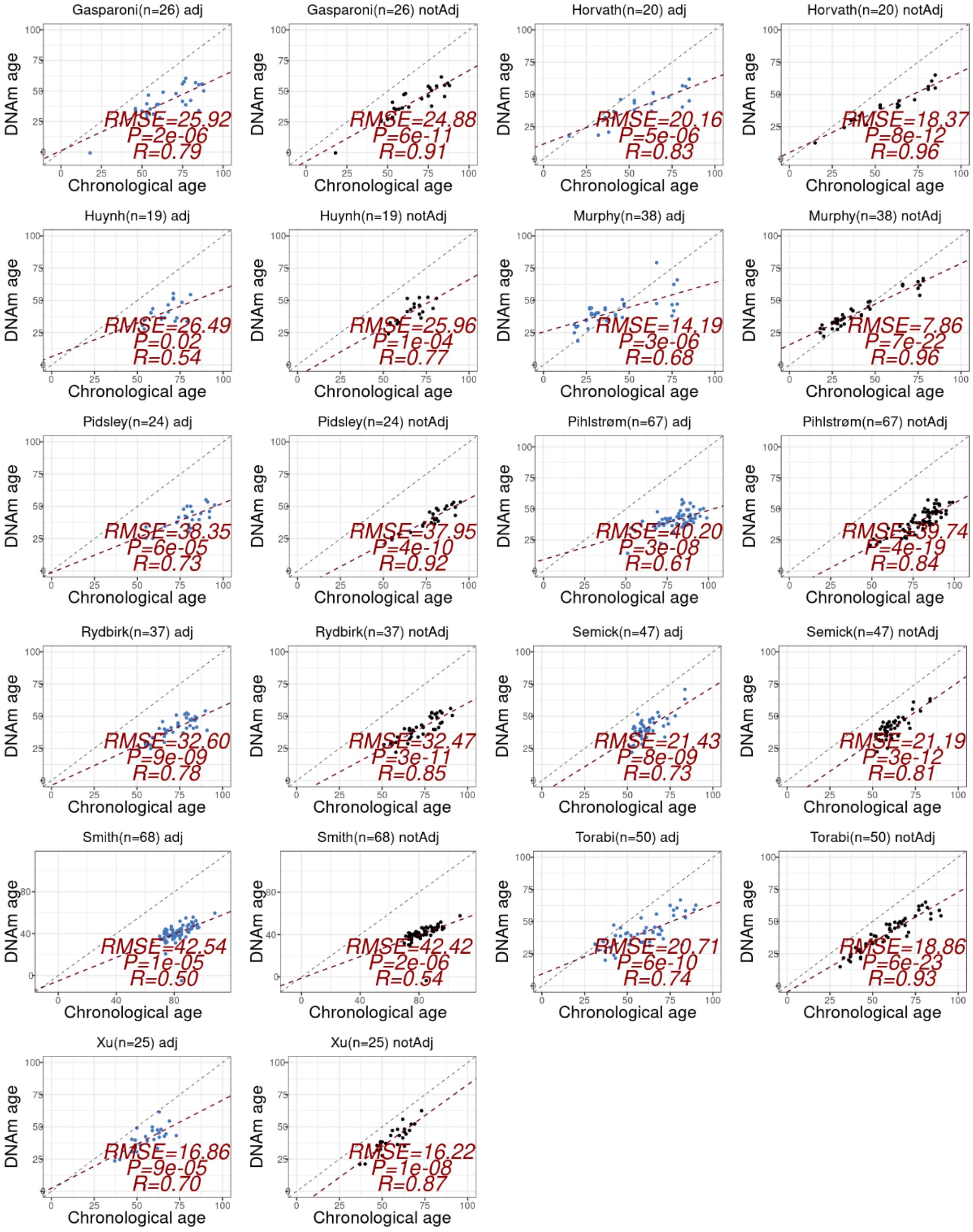


**SI fig.S3: Predicted DNAmAge vs chronological age for CTH adjusted (7 brain cell types) and unadjusted clocks in 11 brain-tissue cohorts.** Each scatterplot is labeled by cohort, cohort-size and whether it is adjusted or unadjusted clock. Root Mean Square Error (RMSE), two-tailed P-value of a linear regression and R (Pearson Correlation Coefficient) value is given. Adjusted clock was adjusted for fractions of excitatory and inhibitory neurons, astrocytes, endothelial cells, microglia, oligodendrocytes+oligodendrocyte progenitor cells and other stromal cells.

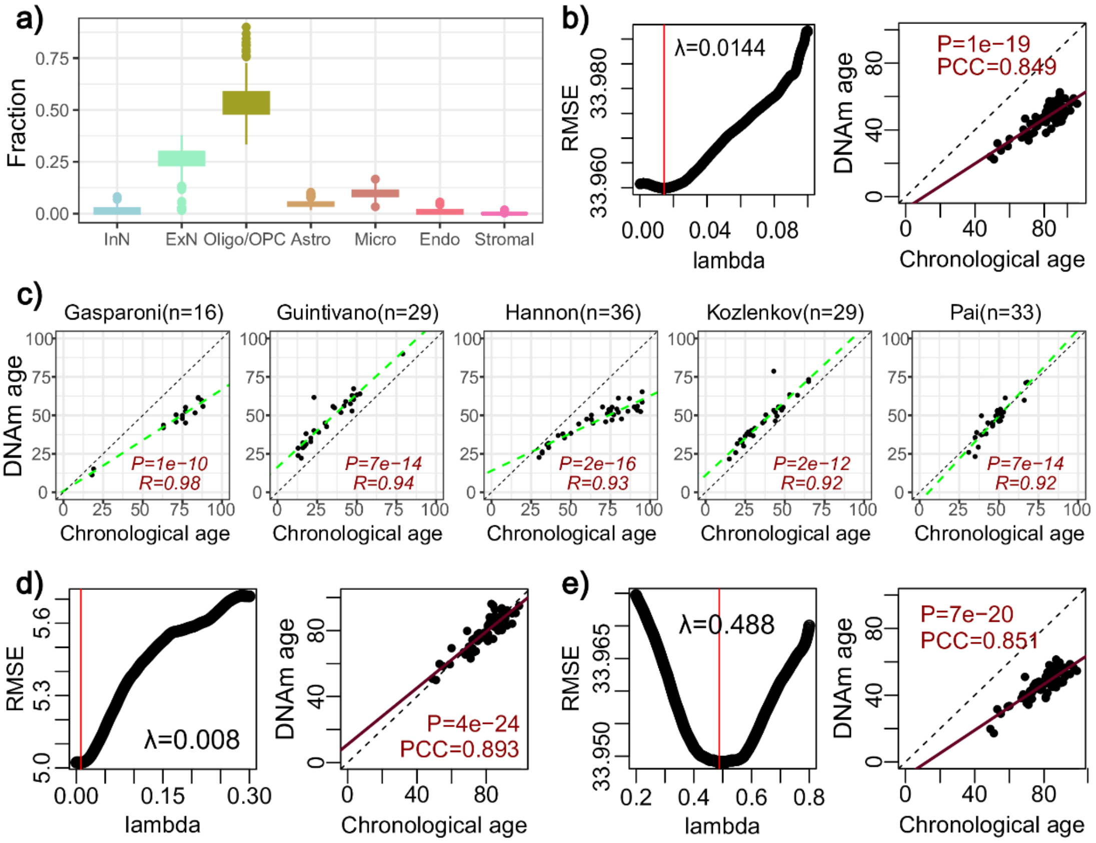


**SI fig.S4: Construction and validation of Neu-In, Neu-Sin and brain-clocks. a)** Boxplot shows the estimated fractions of 7 brain cell types in the Jaffe cohort, as estimated using HiBED. InN: inhibitory neuron; ExN: excitatory neuron; Oligo/OPC: oligodendrocyte and oligodendrocyte precursor cell; Micro: microgliacyte; Endo: endothelial cell; Stromal: stromal cell. **b)** Optimization and validation of Neu-In clock. Panel on the left shows how RMSE changes with the penalty parameter lambda. Red vertical line represents the optimal lambda value. Panel on the right shows the validation of the optimal clock in Philstrom cohort. PCC-value and associated two-tailed P-value are given. **c)** Scatterplots of the chronological age (x-axis) vs predicted DNAm-age (y-axis) from the Neu-In clock in 5 sorted neuron cohorts. Cohort sizes are indicated above each plot. R-value and associated two-tailed P-value from a linear regression test are given. **d)** Same as b), but for the Neu-Sin clock. **e)** As b) but for the BrainClock.

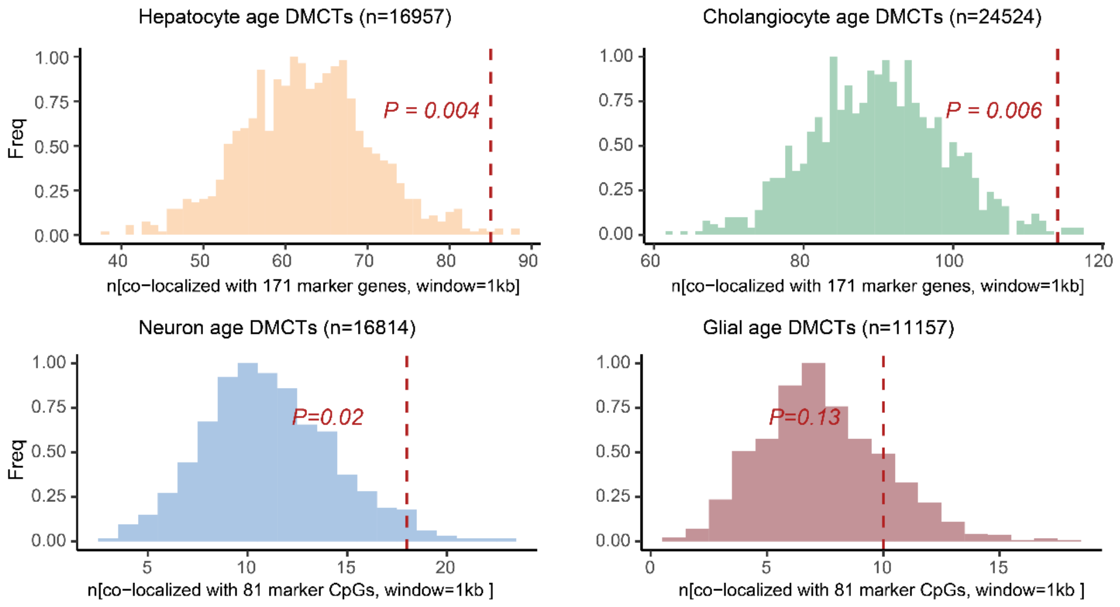


**SI fig.S5: Co-localization of age-DMCTs with DNAm reference matrix CpGs.** For four classes of age-DMCTs (hepatocyte, cholangiocyte for liver and neuron, glia for brain) we display their overlap with the CpGs in the liver and brain DNAm reference matrices, respectively. We used a 1kb window, so that if an age-DMCT is within 1kb of a corresponding marker CpG in the DNAm reference matrix, this was counted as a “hit”. We then compared the observed overlap (red vertical dashed line) with the overlap expected had we chosen an exact same number of “age-DMCTs” randomly from the array. The latter randomization was done 1000 times, yielding a null distribution that we can compare to, to hence derive an empirical P-value.

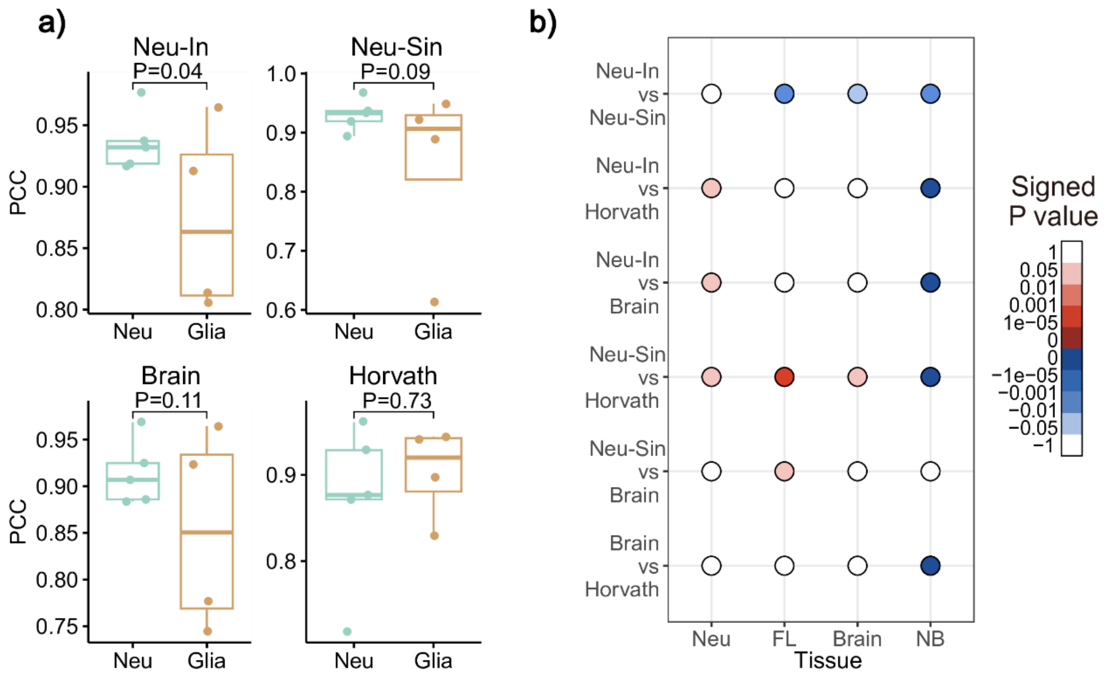


**SI fig.S6: Specificity of neuron-clocks. a)** Neuron specific clocks show specificity to neurons according to a weighted linear regression of PCC values against cell type. Boxplots compare PCC values between predicted age and chronological age (y-axis) in cohorts from 2 cell types (x-axis) for the Neu-In clock, Neu-Sin clock, Brain clock and Horvath clock. There are 5 sorted neuron cohorts and 4 glia cell cohorts in total. P values are from a linear regression of PCC against cell-type (Neu vs Glia) weighted by cohort size. **b)** Comparison of Neu-In, Neu-Sin, Brain and Horvath clock in sorted neuron cohorts (Neu), bulk frontal lobe cohorts (FL), all brain tissue cohorts (brain) and non-brain-tissue cohorts (NB). Balloon plot shows signed P-values from a one-tailed paired Wilcoxon test comparing PCC values of one clock to another. PCC-values represent the correlation between predicted DNAm-Age and chronological age, as obtained with a given clock. Because there are 4 clocks (Neu-In clock, Neu-Sin clock, Brain clock and Horvath clock), there are 6 pairwise comparisons (y-axis). x-axis labels the datasets over which the P-values were estimated. Positive signed P-values indicate higher PCC values for the first clock in the pairwise comparison (e.g. for “Neu-In vs Neu-Sin”, a positive signed P-value means PCC values of Neu-In are bigger than Neu-Sin).

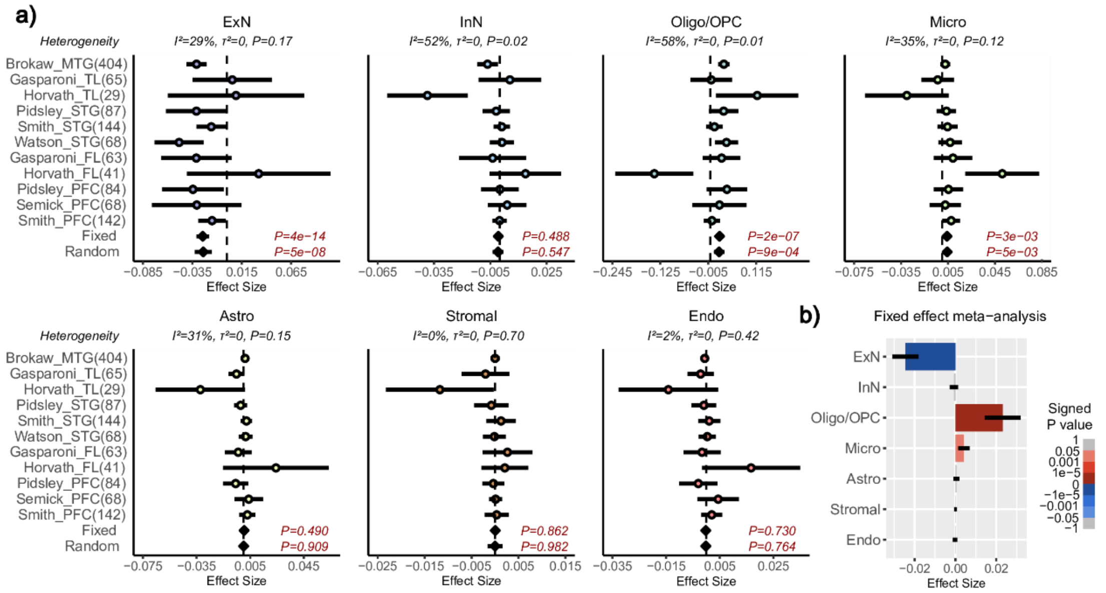


**SI fig.S7: Meta-analysis of brain cell-type fraction associations with Alzheimer's Disease (AD). a)** Forest plots of associations between cell-type fractions and AD, for 7 brain cell-types (ExN=excitatory neurons, InN=inhibitory neurons, Oligo/OPC=oligodendrocytes/oligo precursor cells, Micro=microglia, Astro=astrocytes, Stromal, Endo=endothelial) across 11 independent DNAm datasets. Cohorts are labeled by name and brain region. Brain regions: Middle and Superior temporal gyrus (MTG and STG), temporal lobe (TL), frontal lobe (FL) and prefrontal cortex (PFC). Number of samples in each cohort is given in brackets. P-values from a Fixed and Random Effects models are also given. Heterogeneity statistics and P-values of heterogeneity are given above each plot. **b)** Effect sizes and P-values from the Fixed Effects meta-analysis model.

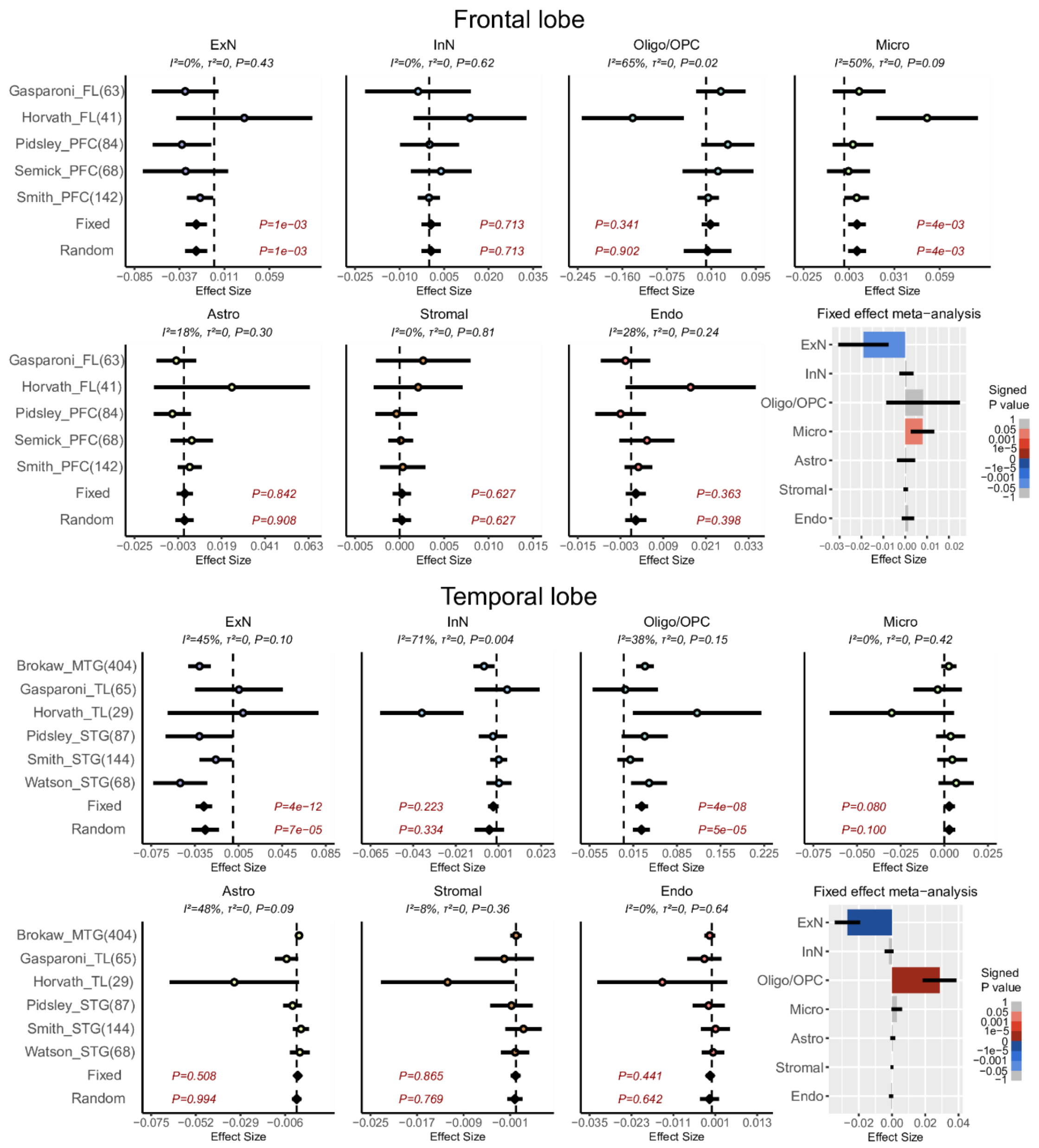


**SI fig.S8: Meta-analysis of brain cell-type fraction associations with Alzheimer's Disease (AD) stratified by brain region.** Top panel: Forest plots of associations between cell-type fractions and AD, for 7 brain cell-types (ExN=excitatory neurons, InN=inhibitory neurons, Oligo/OPC=oligodendrocytes/oligo precursor cells, Micro=microglia, Astro=astrocytes, Stromal, Endo=endothelial) across 6 independent frontal lobe DNAm datasets. Cohorts are labeled by name and brain region/subregion. Brain regions: Middle and Superior temporal gyrus (MTG and STG), temporal lobe (TL), frontal lobe (FL) and prefrontal cortex (PFC). Number of samples in each cohort is given in brackets. P-values from a Fixed and Random Effects models are also given. Heterogeneity statistics and P-values of heterogeneity are given above each plot. Effect sizes and P-values from the Fixed Effects meta-analysis model are given separately. Bottom panel: as top, but for a meta-analysis over 5 independent temporal lobe DNAm datasets.

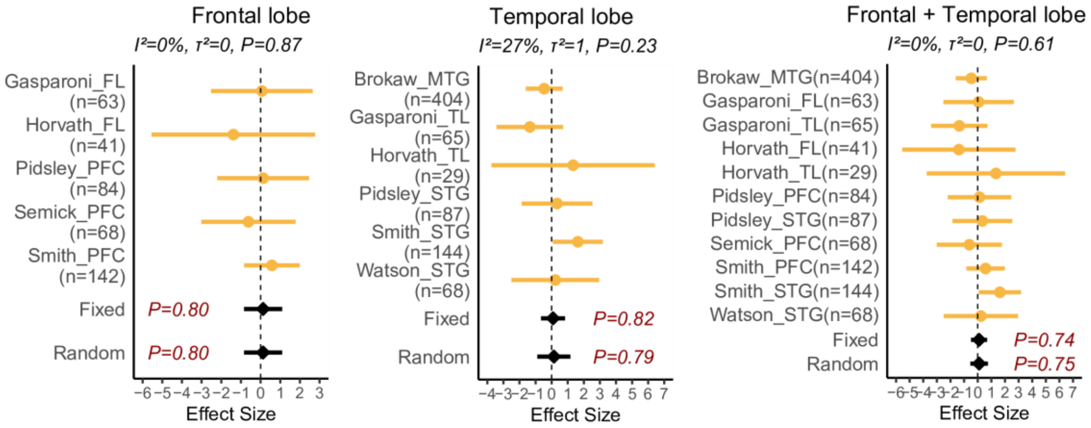


**SI fig.S9: Meta-analysis of Horvath clock associations with Alzheimer's Disease (AD) stratified by brain region.** Top panel: Forest plots of associations of Horvath's clock age-acceleration adjusted for age, sex and brain cell-type fractions with AD. Number of samples in each study is given in brackets. P-value of heterogeneity is given above each panel. P-values of a fixed and random effect models are given.

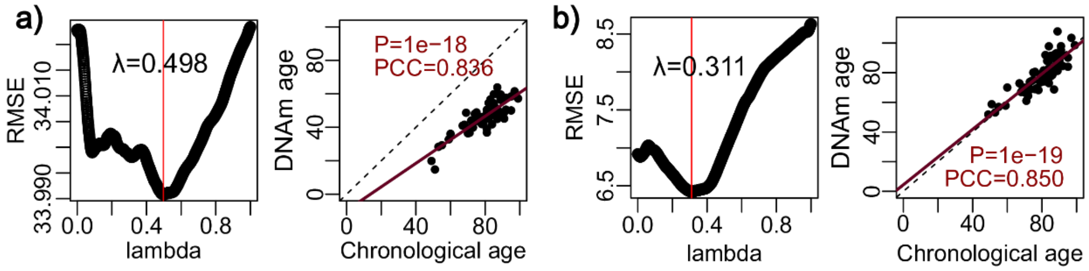


**SI fig.S10: Optimization and validation of Glia-In and Glia-Sin clocks. a)** Panel on the left shows how RMSE changes with the penalty parameter lambda for the Glia-In clock. Red vertical line represents the optimal lambda value. Panel on the right shows the validation of the optimal clock in Philstrom cohort. PCC-value and associated linear regression two-tailed P-value are given. **b)** As a) but for the glia semi-intrinsic (Glia-Sin) clock.

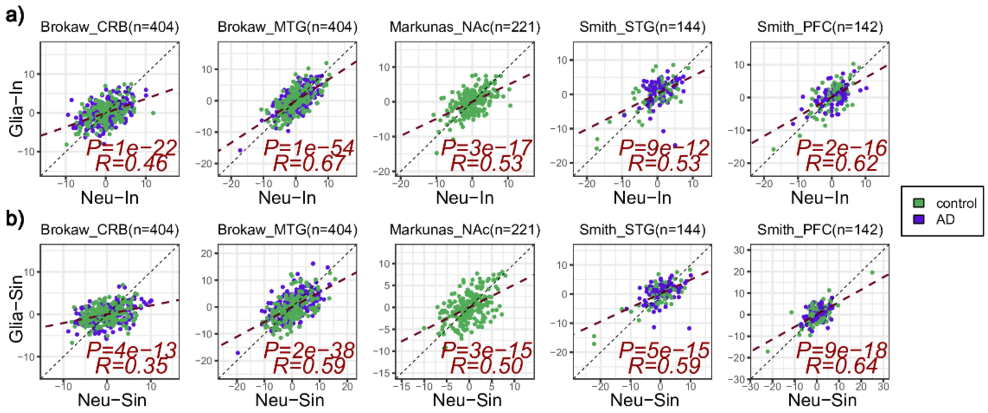


**SI fig.S11: Correlation of neuron with glia clocks in AD cohorts. a)** For the 5 largest brain-tissue cohorts with AD cases and controls, we display a scatterplot of the age-acceleration (extrinsic age acceleration-EAA) of the Glia-In clock (y-axis) vs the age-acceleration (EAA) of the Neu-In clock (x-axis). The R-values and two-tailed P-values from a linear regression are given. The number of samples in cohort is indicated above the plot. **b)** As a) but for the Glia-Sin and Neu-Sin clocks. In the two Smith datasets and for both a) and b) an outlier control sample with very negative age-acceleration has been capped at a value given by the minimum across all other samples.

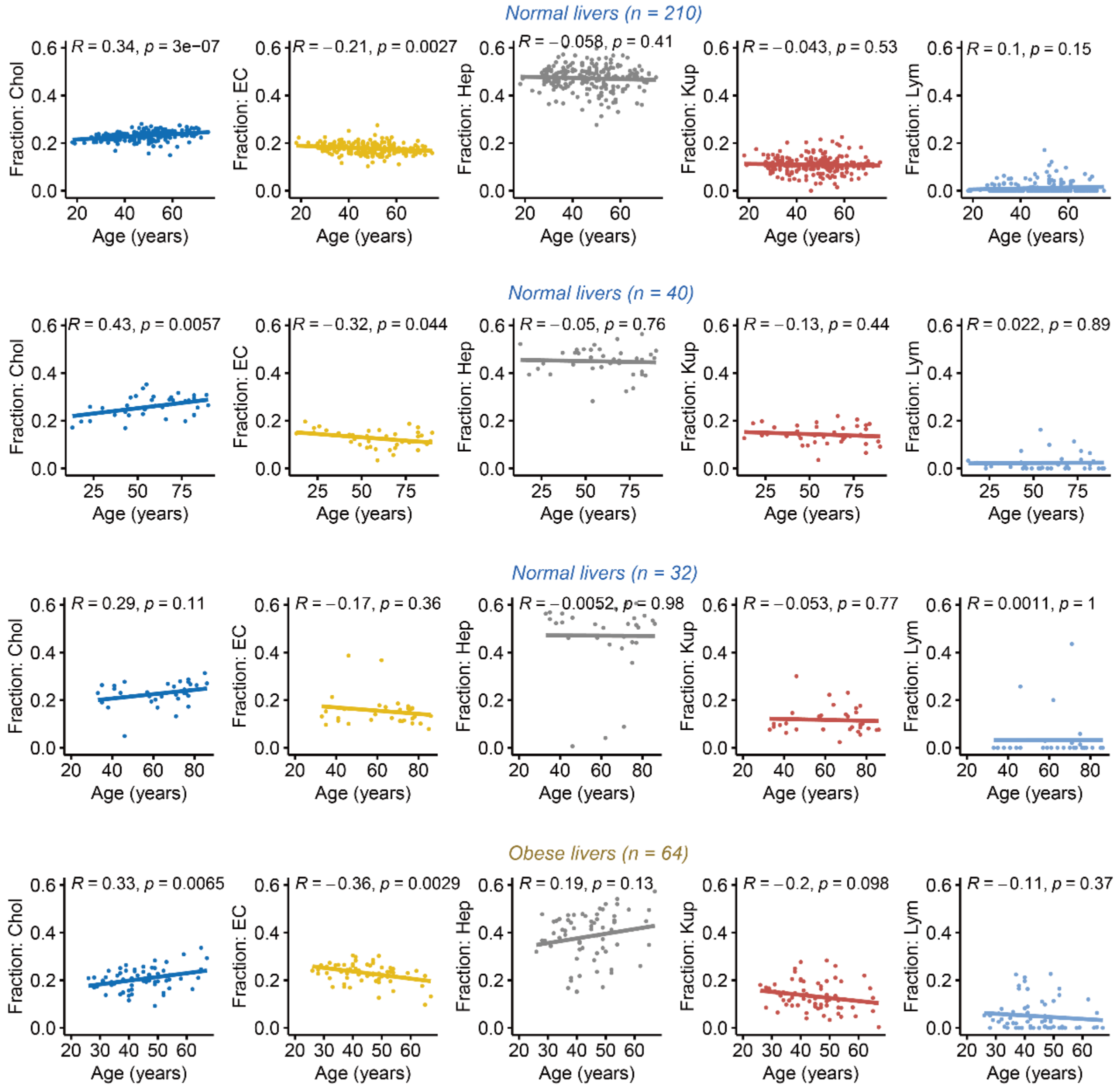


**SI fig.S12: Liver cell-type fraction variations with age.** For four independent DNAm liver datasets, we plot the fractions of five liver cell-types (as estimated using EpiSCORE) as a function of age. R-values and P-values (two-tailed linear regression) of statistical significance are given. From left to right, cell-types are Cholangiocytes (Chol), endothelial cells (EC), Hepatocytes (Hep), Kupffer macrophages (Kup) and Lymphocytes (Lym).

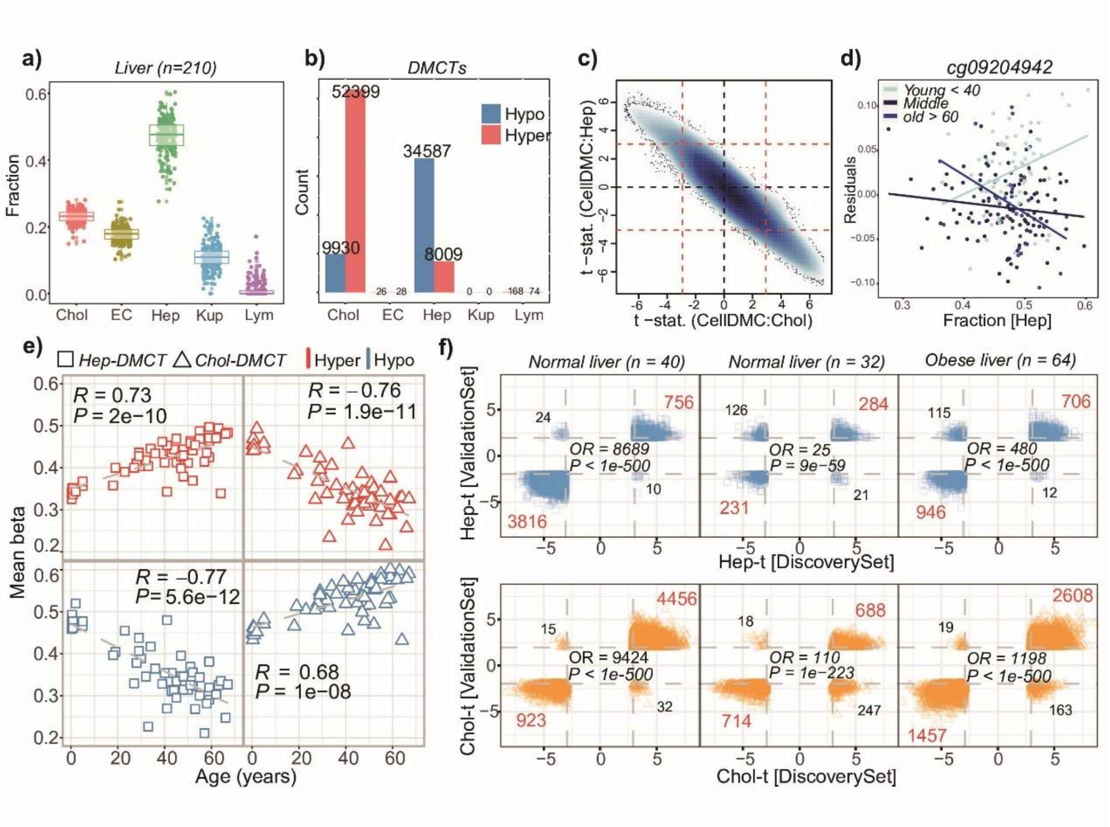


**SI fig.S13: Identification and validation of hepatocyte-specific age-DMCs. a)** Boxplots of estimated cell-type fractions (y-axis) using EpiSCORE's DNAm liver reference matrix of 5 liver cell-types (Chol=cholangiocytes, EC=endothelial cells, Hep=hepatocytes, Kup=Kuppfer macrophages, Lym=lymphocytes, x-axis) in a series of 210 normal liver samples. **b)** Barplots displaying the number of hypo and hypermethylated age-DMCTs for each cell-type, as inferred with CellDMC using the estimated fractions in a). **c)** Smoothed scatterplot of CellDMC t-statistics for age-DMCTs in hepatocytes (y-axis) vs cholangiocytes (x-axis). Red dashed lines indicate the FDR=0.05 significance level. **d)** Example of a hepatocyte-specific age-DMCT with y-axis labeling the residual after adjusting for cell-type fractions and x-axis labeling the hepatocyte fraction of the sample, with samples colored by age-group. **e)** Scatterplot of average DNAm levels (Mean Beta, y-axis) of four sets of age-associated DMCTs against chronological age (x-axis) in an independent DNAm dataset comprising 55 hepatocyte cultures. The four sets of CpGs are age-associated hypermethylated and hypomethylated DMCs in hepatocytes and cholangiocytes, respectively. In each panel, we give the Pearson Correlation Coefficient (PCC) and associated correlation-test P-value. **f)** Top row: Scatterplots of CellDMC t-statistics for hepatocyte-specific age-DMCTs in the discovery set (x-axis) vs validation set (y-axis) for 3 separate validation sets encompassing bulk liver tissue samples, as shown. Red dashed lines indicate FDR=0.05 (discovery set) and P=0.05 (validation set). In each panel, we give the number of age-DMCTs that fall in each significant quadrant, and report the Odds Ratio (OR) and associated P-value from a one-tailed Fisher-test. Bottom row: as top row but for cholangiocyte-specific age-DMCTs.

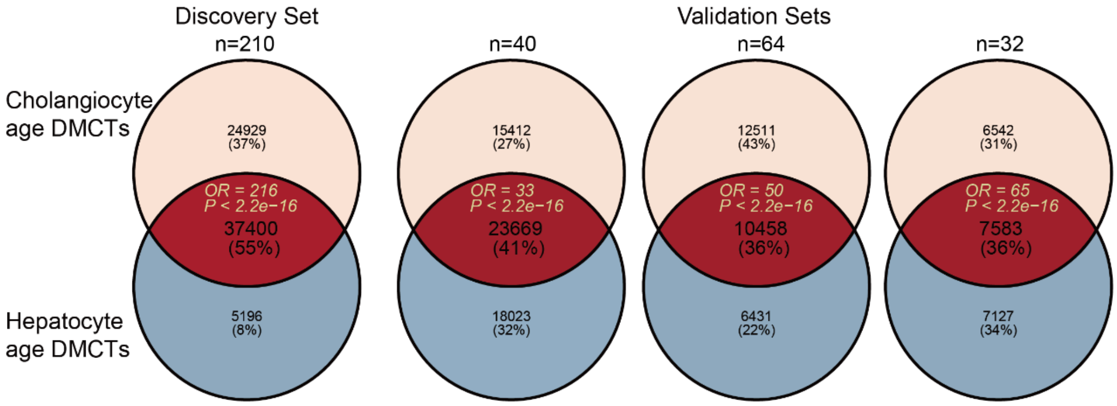


**SI fig.S14: Venn Diagrams of overlap between hepatocyte and cholangiocyte age-DMCTs.** For the discovery liver DNAm dataset, as well as 3 independent validation liver sets, we display Venn Diagrams showing the overlap of hepatocyte and cholangiocyte age-DMCTs, as inferred using CellDMC. Odds Ratios (OR) and P-values of overlap were obtained from one-tailed Fisher exact tests.

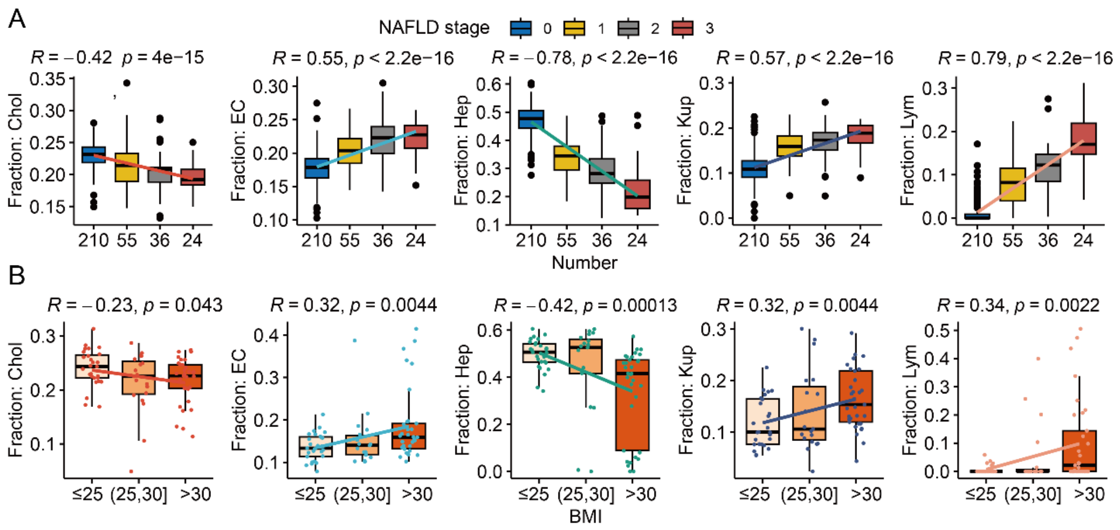


**SI fig.S15: Variations of liver cell-type fractions with NAFLD and obesity. A)** Boxplots display the estimated fraction of a given cell-type as a function of NAFLD stage (0=normal) for cholangiocytes (Chol), endothelial cells (EC), hepatocytes (Hep), Kupffer macrophages (Kup) and lymphocytes (Lym). R-values and two tailed P-value of a linear regression are given. Number of samples in each stage is given below x-axis. **B)** As A) but with the boxplots displaying the fractions as a function of body mass index (BMI) with BMI stratified into 3 categories, as shown. Of note, A) and B) are for independent DNAm datasets, as described in Methods.

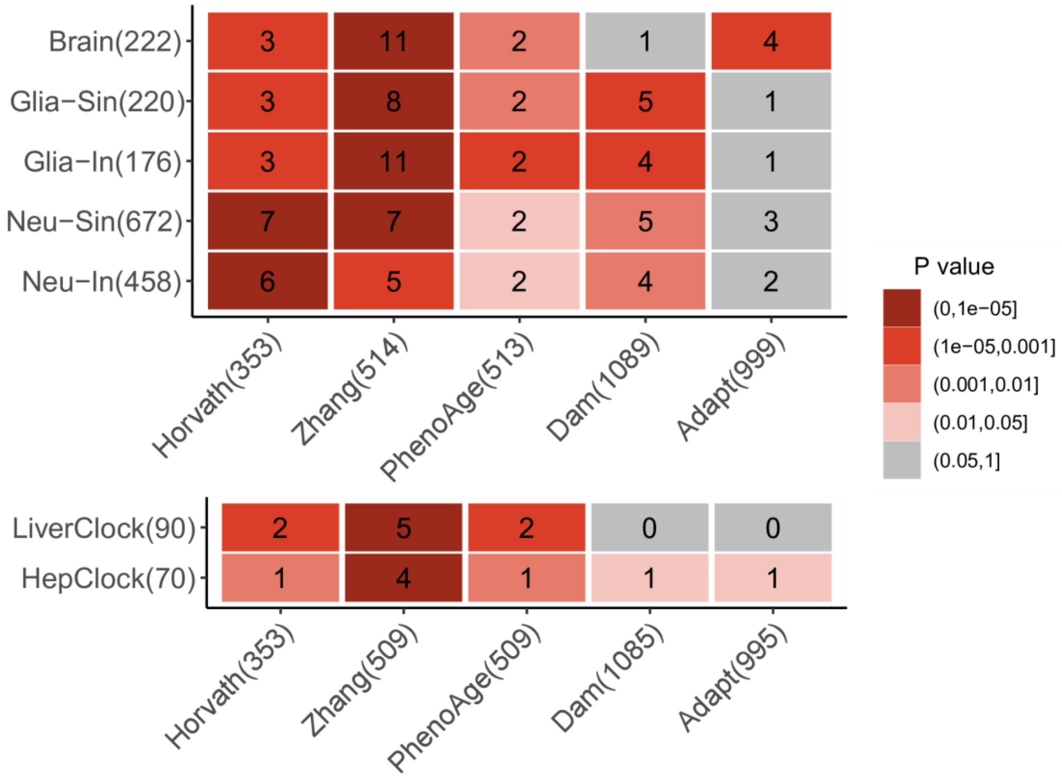


**SI fig.S16: CpG overlap diagram of cell-type specific clocks with other clocks.** For 5 cell-type specific clocks as well as 2 tissue-specific ones (Brain & Liver), depicted on the y-axis, we display the overlapping CpG number as well as their statistical significance, with the CpGs making up well-known epigenetic clocks (x-axis). The P-values derive from a one-tailed binomial test. The number in brackets is the number of clock CpGs present in the DNAm dataset.

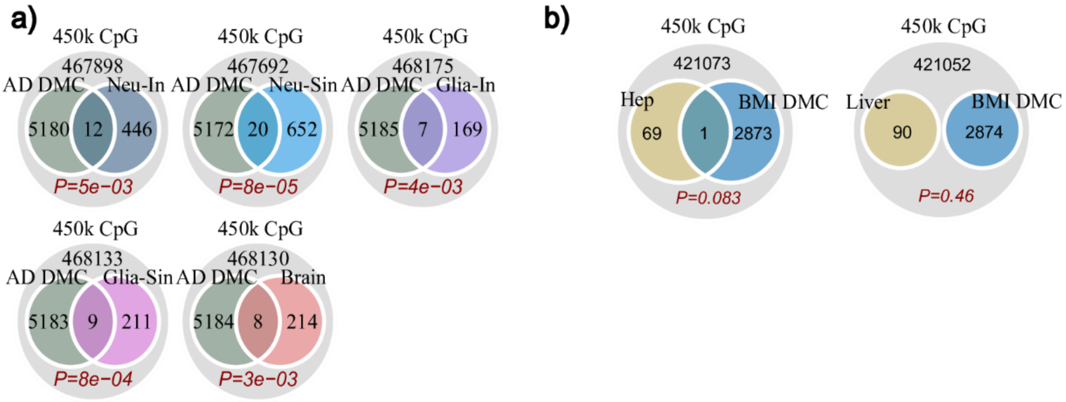


**SI fig.S17: Enrichment analysis of clock-CpGs for EWAS DMCs. a)** Venn diagrams displaying the overlaps of 5 brain-related clocks with Alzheimer's Disease (AD) associated DMCs, with the one-tailed P-values derived from a Binomial-test. **b)** As a) but for the Hep and Liver clocks.

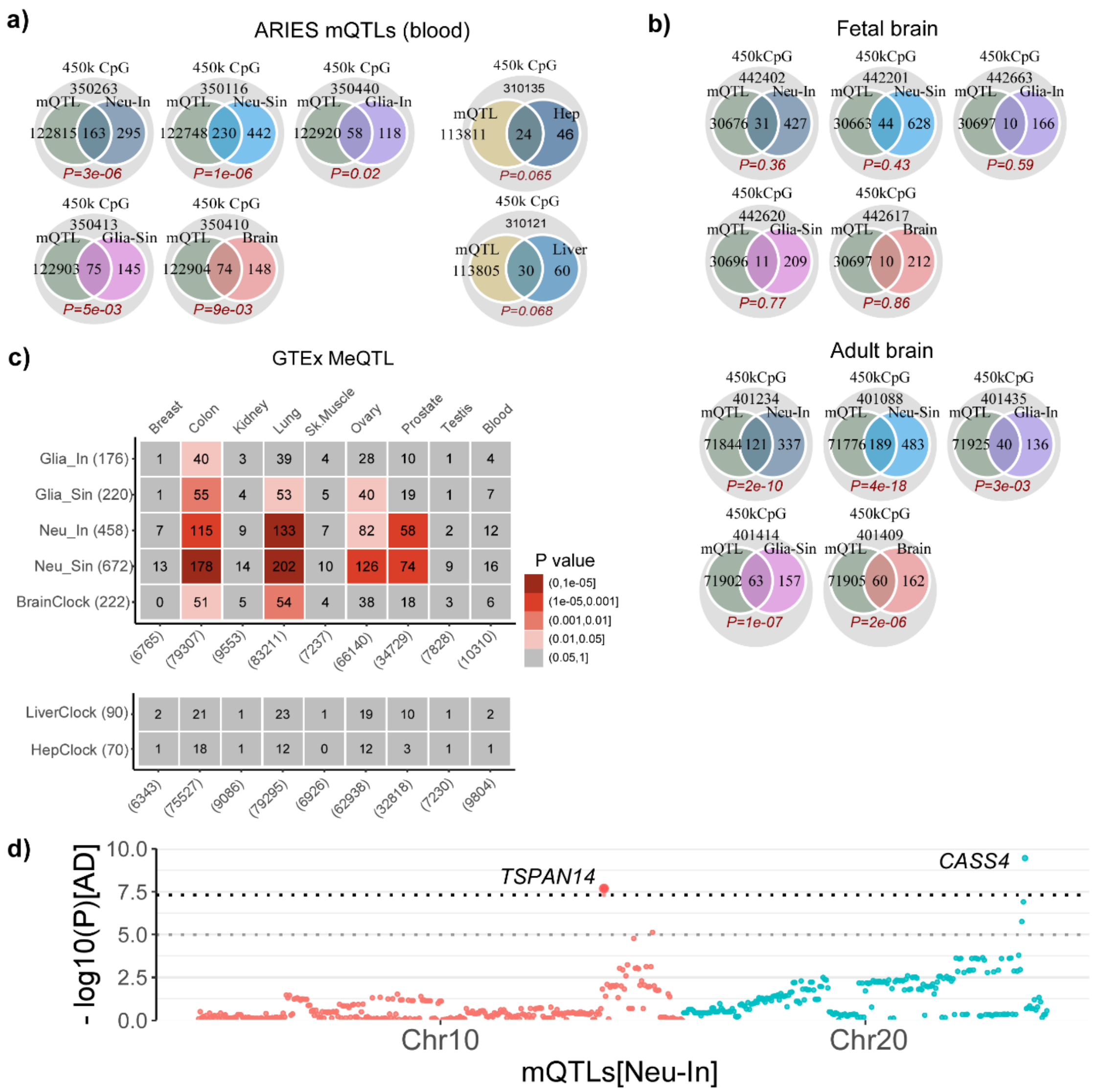


**SI fig.S18: Enrichment analysis of clock-CpGs mQTLs. a)** Venn diagrams displaying the overlaps of 5 brain-related and 2 liver-related clocks with blood mQTLs as defined by the ARIES database. P-values computed using a one-tailed Binomial test **b)** As a) but only for brain-related clocks and mQTLs as defined from fetal and adult brain. **c)** As a) but for mQTLs as defined in various tissue-types from eGTEX. The numbers in brackets indicate the number of clock-CpGs or the number of distinct mQTLs (CpGs) of each tissue present in the corresponding DNAm dataset. **d)** Manhattan like plot of two Neu-In clock CpGs that map to cis-mQTLs in blood and adult brain and where the associated gene has been previously implicated in Alzheimer's Disease (AD). Y-axis labels the significance levels of the SNP associations with AD.
